# LUC3D: Multi-animal 3D pose tracking in the browser

**DOI:** 10.64898/2026.11.17.767188

**Authors:** Eric J. Leonardis, Stefan N. Oline, Isaac Tang, Joshua Park, Bartul Mimica, Liezl Maree, Shayan Afshar, Olivia Timmermans, Jinrun Jiang, Eliza Whitmire, Aaron Zhu, Talisa Wines, Kay M. Tye, Mala Murthy, Annegret L. Falkner, Talmo D. Pereira

## Abstract

Animal behavior unfolds in three dimensions over time, yet accessible scientific tools for analyzing it in its natural form remain limited. Uncertainty from popular 2D pose methods can be refined with multi-camera approaches to produce more accurate 3D estimates. This paper presents LUC3D (Label Unification and Correspondence in 3D Annotation GUI), a browser-based 3D GUI for reducing the amount of manual labor required for labeling, proofreading, and multi-camera multi-animal tracking while requiring no installation to maximize accessibility. As proof-of-concept we present two datasets of proofread multi-camera 3D triangulated pose data of mouse social behavior, a protocol for generating 3D consistent labels, and a cross-view re-identification tracking algorithm for multi-animal multi-camera pose data. This tool allows for the widespread adoption of 3D pose estimation techniques for studying social behavior while reducing the cost of entry for neuroscience labs across the world.

## 1 Introduction

Markerless pose estimation has transformed the quantification of animal behavior [1–4]. Single-camera methods track keypoints in 2D pixel coordinates, so they cannot reliably measure metric distances or recover the vertical axis. The human pose estimation literature has demonstrated that the solution to the limitations of 2D is to use multiple camera views to enable 3D pose estimation [5]. This methodology has been applied to 3D pose estimation of animal behavior [6–8], and Anipose in particular has become a widely used and reliable tool for camera calibration and triangulation [7]. These single-subject 3D methods have since been extended to multiple individuals [9, 10] and to multiple animals [11–13]. Dedicated tools for 3D annotation have also been developed [14, 15]. These tools are effective and widely used, and the remaining gaps lie in their speed, their accessibility and how they fit together. Markerless predictions also still need human proofreading. In rats, where markerless keypoint trackers have been compared against skin-attached markers, their predictions were precise to about ±10 mm, comparable to the distance between many landmarks on the mouse [16, 17]. Multiple animals add a further challenge that single-animal 3D methods never face. Before any keypoint can be triangulated, the detections of each animal must be matched across every camera. A wrong match blends two animals into one 3D pose, and the error then appears in every view. Resolving cross-view identity matching remains a central challenge in 3D multi-animal pose estimation [10, 18].

Despite this progress, 3D multi-animal pose estimation remains challenging to adopt. It is still out of reach for labs without the resources to install and maintain these tools. Most require a local installation and are built around a GPU workstation which limits adoption in labs without dedicated engineering support [14]. Yet labeling and proofreading make up much of the work in a 3D project, and neither needs a GPU. The workflow is also fragmented across separate packages for calibration, labeling, cross-view identification and proofreading. Labeling also scales with the number of cameras. Single-camera pose estimation methods require many manually labeled frames per view, so every camera added to a rig adds another set of frames to label. Identity must be resolved multiple times. Each animal must be tracked over time within every view and then matched across views, which has been done by manual assignment, color marking or the assumption that the initial tracks have no dropped frames [13, 19]. Usually, proofreading identity has to be repeated once per camera which can be laborious. Progress is further limited by the lack of large benchmark datasets of multiple animals in enriched environments with heavy occlusion.

An alternative is to move every step after inference into the browser and to resolve identity with 3D geometry. Such a tool would run on any computer, with no installation and no dedicated GPU. Labelers and proofreaders could then work in parallel within a lab’s existing SLEAP workflow while models are trained and run on one GPU machine [3]. It would use multi-view geometry to reduce manual work. Triangulation recovers a 3D point from its 2D detections in multiple views, and reprojection projects that point back into each camera, so labels placed in a few views can fill in the rest. The reprojection error can be calculated by the pixel distance between the reprojected points and the original labels before triangulation and used to inform the user about agreement between cameras. This can also be used to resolve identity across all cameras at once from 3D geometry, without physically marking the animals [10, 18]. Triangulation and tracking also need to be fast enough for interactive use.

Here we present LUC3D, a browser-based GUI for 3D multi-animal labeling, cross-view tracking, triangulation and proofreading. First, we release two benchmark datasets. MouseDyad-10M contains 55 sessions of male-female mouse pairs, with 49.5 million video frames, 19.8 million 3D poses triangulated from identity-proofread cameras and 99.0 million 2D poses reprojected from the 3D. SLAP-2M contains 74 sessions of one to four mice across graded levels of environmental enrichment, with 15.6 million video frames and 2.81 million 3D poses. Its sessions span a range of difficulty set by the number of objects and animals, and across all 74 sessions 95% of keypoints are seen by at least two of the six analyzed cameras, so they can be triangulated and reprojected into the views that miss them. Second, we show that a greedy cross-view tracker with a 3D correspondence term resolves identity across all cameras at up to 14,694 frames per second on a laptop. On MouseDyad-10M it keeps identity switches below one per 100,000 camera frames and reaches a mean cross-view IDF1 of 0.971. Over the multi-animal SLAP-2M sessions its mean cross-view IDF1 is 0.932. On SLAP-2M the median is 0.998 for two mice without objects and falls to 0.84 in the most enriched sessions. The same tracker also recovers identity from 3D geometry alone in rats [13], with a mean cross-view IDF1 of 0.985 over 41 sessions. Third, we show that reprojection-aided labeling cuts manual placements in half on a five-camera rig, with savings that grow with every camera added. With five cameras, LUC3D’s 3D poses land a median of 1.2 mm from the dataset’s reference reconstruction, and it triangulates up to 16,030 frames per second. Finally, we use LUC3D to characterize a female-led display during male-female social interactions based on its distinctly 3D kinematics. Together with a browser-based calibration tool, LUC3D takes a lab from calibration to proofread 3D poses of interacting animals without an installation or a GPU on every lab computer. LUC3D is accessible at https://luc3d.sleap.ai.

## 2 Results

### 2.1 LUC3D resolves cross-view identity and 3D pose in the browser

Multi-camera pose recording introduces the problem of linking identities across views, in addition to the single-camera problem of linking them across frames. With several animals, a 2D tracker in each camera links each animal’s detections from frame to frame into a track, however a track exists only within its own camera and its name is arbitrary, so the same mouse often belongs to a different track in each view (Figure 1a, *top*). Triangulation instead needs a single identity per animal that is shared by all of the cameras (Figure 1a, *middle*). Existing detectors and proofreading tools produce these per-camera tracks, but there is no standard tool that links them into identities, leading to a large manual proofreading load for the user that scales linearly as the number of cameras increases. LUC3D links identities automatically by matching, in every frame, each camera’s detections to the animals’ current positions in 3D, so the tracks that follow the same animal in different cameras are given the same identity. On any given frame from an eight-camera recording of three mice, the detections carry 21 different track names across the views, which LUC3D resolves into three identities, one per animal. Importantly for an automated process, it also leaves out one duplicate detection of an animal that it has already matched.

**Fig. 1:**
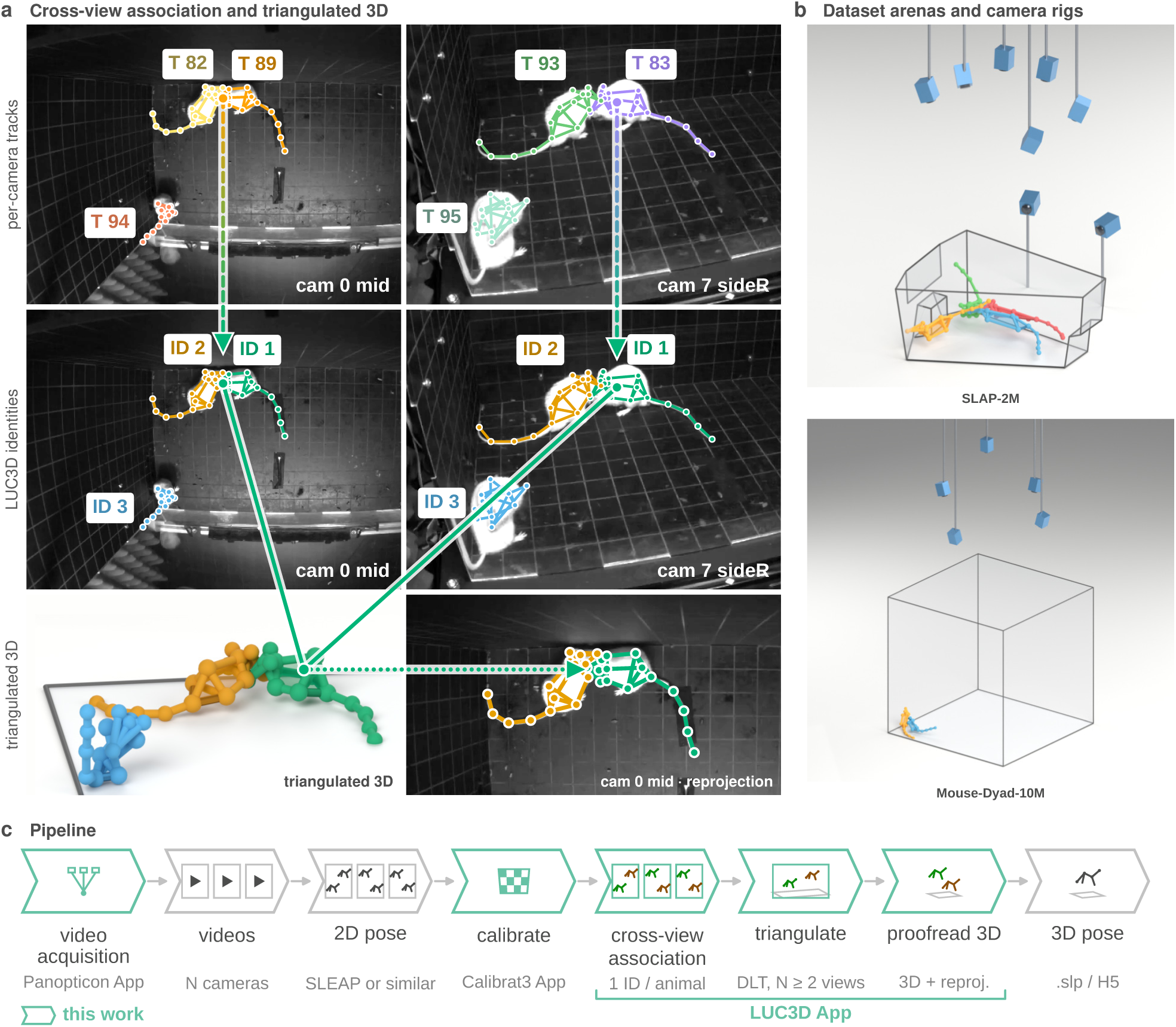
LUC3D enables annotation and proofreading multi-camera 3D pose in a browser with no installation. **a**, Cross-view association and triangulation on one frame of an 8-camera, 3-mouse recording with a 15-node skeleton. The two columns are the same two views, cam 0 mid and cam 7 sideR. Top, the per-camera tracks supplied to the app, each labeled with the track name its own camera assigned. Track colors are arbitrary, so the same animal is a different color and a different track name in the two views. Middle, the same two views after association, each labeled with the resolved identity. Identity colors are shared by every view, so each animal keeps one color and one number in both views. Bottom left, a render of the triangulated 3D poses. Bottom right, the 3D poses reprojected into cam 0 mid, magnified. The lines follow one animal, ID 1. Dashed arrows point from its two tracks to its identity, solid lines join its identity in each view to its 3D pose, and the dotted arrow points to its reprojection. **b**, The two datasets’ arenas with their camera rigs and poses. Top, the SLAP-2M rig with 8 calibrated cameras and the cage with the tracked 3D poses of four animals. Bottom, the MouseDyad-10M rig with the 5 calibrated cameras and enclosure with tracked 3D poses of the two animals. **c**, Pipeline of a standard recording and labeling session. Videos from N synchronized cameras are acquired with Panopticon. A 2D pose model (SLEAP or any other predictor, as .slp) predicts keypoints in every view, and the cameras are calibrated with Calibrat3, a browser-based tool. The LUC3D app then associates detections across views into one identity per animal, triangulates them from two or more views, and proofreads the 3D poses against per-view reprojections. The output is a 3D pose for each animal, saved as .slp or HDF5. Green outlines mark the stages contributed by this work.

To test LUC3D’s robustness, we evaluated its performance on the two large provided datasets spanning a range of technical and experimental conditions (Figure 1b). SLAP-2M contains 74 sessions and 15.6 million video frames, recorded with eight cameras, of one to four mice among graded levels of environmental enrichment. MouseDyad-10M contains 55 sessions and 49.5 million video frames, recorded with five cameras, of male-female pairs. Together they vary the number of cameras, the number of animals and how often objects or other animals occlude an animal from view. Supplementary Videos 1 and 2 show the two rigs and their tracked 3D poses in motion. By bringing cross-view identity, support for multiple animals and a 3D proof-reading view together in one browser app with no installation, LUC3D makes 3D multi-animal labeling more easily accessible than existing tools [14, 20].

LUC3D is one part of a pipeline that runs from recording to calibration to proofread 3D poses (Figure 1c). Videos are first recorded from hardware-synchronized cameras. We provide Panopticon, an acquisition app that records trigger-synchronized video from every camera and can calibrate the rig from the same window (Supplementary Figure 1). The user acquires a brief video of a ChArUco board in a variety of positions, and the cameras are then calibrated from this session with calibrat3, which runs in the browser (Supplementary Figure 2). Next, a 2D pose model such as SLEAP predicts keypoints in every view [3], since LUC3D does not detect keypoints itself. The user then uses LUC3D to load those predictions, assign cross-view identities, triangulate the keypoints into 3D, and proofread the 3D poses against every camera’s video, all in a web browser with no installation (Supplementary Figure 3). Projects are saved as SLP files, so they open in SLEAP as well as in LUC3D. The output is a 3D pose for each animal, with one identity shared across all cameras. Within each frame, LUC3D triangulates all 45 keypoints of the three mice, and each 3D pose can be overlaid (reprojected) onto the video of any camera for visual inspection (Figure 1a, *bottom*).

### 2.2 Greedy cross-view association is fast and accurate

LUC3D compares each detection with a 3D anchor for each animal, the animal’s most recent 3D position (Figure 2a). The comparison has a 2D term in the camera image and a 3D term between the anchor and the camera ray through the detection. Maree et al. (2024) [18] use an exhaustive cross-view association algorithm that enumerates every possible grouping of detections into identities, then triangulates and reprojects each, and keeps the groupings with the lowest reprojection error. Figure 2b shows these hypotheses for three animals in two camera views. Following Chen et al. (2020) [10], LUC3D instead solves one Hungarian assignment per camera, matching that camera’s detections to each animal’s most recent 3D position, and re-triangulates each animal once all cameras have been assigned (Figure 2c). This costs *C* · *A*^3^. Exhaustive enumeration instead requires *A*! hypotheses per view and (*A*!)*^C^*per frame, meaning that there are 32 hypotheses for two animals in five cameras and ∼190 million hypotheses for four animals in six cameras (Figure 2d). Keeping the wrong hypothesis swaps the identities of two animals (Figure 2e). Exhaustive enumeration is undefined unless every camera has exactly as many detections as there are animals, so it was run only on the frames that meet this condition, ∼9.9 million of the ∼10.6 million.

**Fig. 2:**
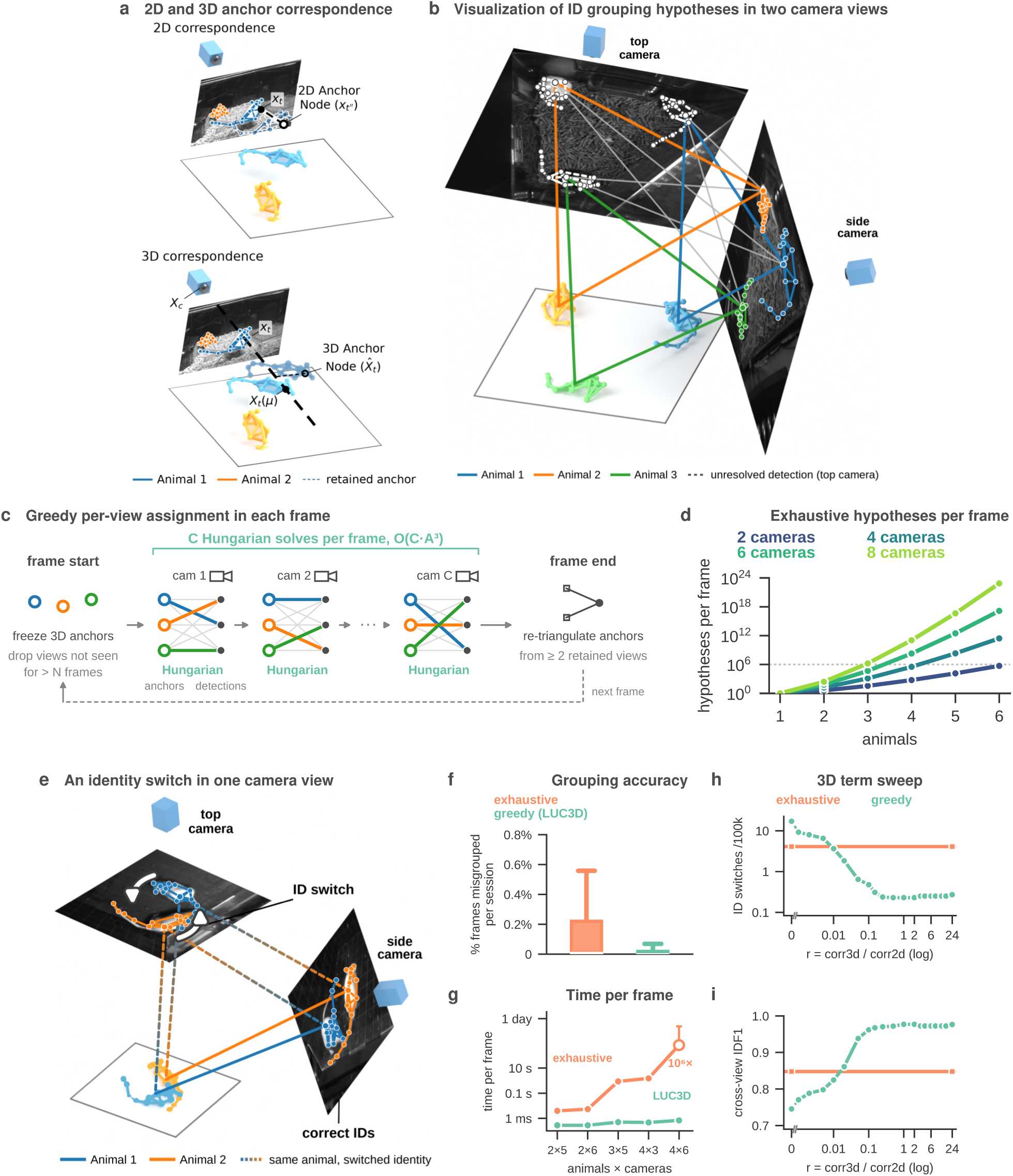
Cross-view association is fast and accurate. **a**, The tracker’s 2D and 3D correspondence terms on one frame. Top, the 2D term (black dashed line) is the image distance between the current detection and that camera’s retained 2D anchor (dashed translucent blue pose). Bottom, the 3D term (dotted black line) is the distance between the 3D anchor node and the ray from the camera through the current node. **b**, The possible groupings for three animals seen by two cameras. Correct groupings are in color and incorrect ones in gray. **c**, LUC3D’s association in one frame. The 3D anchors are frozen at the start of the frame, and detections older than *N* frames are dropped. Each camera’s detections are then assigned to the anchors with the Hungarian algorithm, which grows as *C · A*^3^ for *A* animals and *C* cameras. Gray lines are scored pairings and colored lines kept pairings. Each anchor is re-triangulated once at the end of the frame. **d**, Groupings that exhaustive enumeration [18] must test per frame, which grow as (*A*!)*^C^*. The dotted line marks 10^6^, above which it was not run. **e**, An identity switch. The two identities are exchanged in the top camera and correct in the side camera. A line that changes color marks one animal given two identities. Two arrows indicate the switch between the two instances on the top camera. **f**, Percentage of frames grouped incorrectly in each session, compared with proofread ground truth. Both medians are below 0.01%. A few outlying sessions are not drawn (up to 4.1% for exhaustive enumeration and 3.1% for LUC3D). **g**, Time per frame on the same detections for each combination of animals and cameras. At four animals in six cameras exhaustive enumeration was not run, and the value shown is a lower bound. **h**, Identity switches per 100,000 camera frames as the weight *r* of the 3D term relative to the 2D term is varied, with exhaustive enumeration as a reference. **i**, Cross-view IDF1 over the same range. Panels f and g use the *∼*9.9 million frames of 97 sessions (55 MouseDyad-10M and 42 SLAP-2M) in which every camera holds one detection per animal. Panels h and i use 55 MouseDyad-10M sessions. Panels a and b are frames of SLAP-2M sessions. Box plots show the median and interquartile range, with whiskers at 1.5 times the interquartile range. Frame sampling and timing are described in Section 4.8.

The greedy assignment groups more frames correctly than exhaustive enumeration. Both methods were scored against proofread ground truth on 97 sessions (Figure 2f). These comprised 55 MouseDyad-10M sessions and the 42 multi-animal SLAP-2M sessions. Across ∼9.9 million frames scored by both methods, the greedy assignment grouped every frame correctly in 23 of 97 sessions and exhaustive enumeration in 15 (Figure 2f). Pooled over all frames, the greedy assignment misgrouped 0.019% of frames and exhaustive enumeration 0.027%.

Greedy assignment is also far faster than the exhaustive approach, 14 times at two animals in five cameras. Figure 2g shows the computation time of the two procedures on the same frames. At two animals in five cameras, exhaustive enumeration takes 4.0 ms per frame and the greedy method 0.28 ms. At four animals in six cameras, the greedy method takes 0.69 ms, but exhaustive enumeration exceeds the 10^6^ hypothesis cap and could not be run. The cost shown for that configuration is a lower bound that uses the fastest perhypothesis rate measured anywhere in the sweep. Even at this bound, one second of 30 fps SLAP-2M video would take more than 5 hours to process.

Figures 2h and i examine the contribution of the 3D correspondence term in the association cost (Section 4.8). We swept the ratio *r* of the 3D to the 2D correspondence term from 0 to 24 over the 55 bench-mark MouseDyad-10M sessions. Each setting is scored by two measures. An identity switch is a detection given the identity of a different animal than in the ground truth. Cross-view IDF1 is the share of detections, pooled across all cameras, that are assigned to the correct animal [21]. A score of 1 means every detection is correct (Section 4.12). Exhaustive enumeration can only run on frames that have no missing detections, 47.9 million of the 49.5 million camera frames, and every setting in Figures 2h and i is scored on exactly those frames. Exhaustive enumeration has no 3D correspondence term and switched identity 4.1 times per 100,000 camera frames at a mean cross-view IDF1 of 0.848 over sessions. With the 3D term removed (*r* = 0), the greedy Hungarian association does worse than exhaustive enumeration, with 17.1 switches per 100,000 camera frames and an IDF1 of 0.746. From *r* = 0.02 upward, greedy association does better than exhaustive enumeration on both measures, so the 3D term is what enables the greedy matching to outperform it. The app default, *r* = 6, gives 0.25 switches per 100,000 camera frames at an IDF1 of 0.972 on these frames, 16 times fewer switches than exhaustive enumeration. Over all 49.5 million camera frames, including those with missing detections, its cross-view IDF1 is 0.971 (Supplementary Figure 4a). Every ratio from 3 to 16 gives the same result, so the default sits on the plateau.

Supplementary Figure 4 and Supplementary Sections 5.2, 5.3 and 5.4 compare the LUC3D cross-view tracker with SLEAP and ByteTrack, report the parameter search that reduced identity switches, and test the tracker on rats from the social-DANNCE datasets [13].

### 2.3 Reprojection from two labeled views reduces manual annotation and triangulation is fast

Reprojection-aided labeling cuts the manual work of multi-camera annotation, and the savings grow with every camera added. Traditional manual multi-camera annotation scales linearly with the number of cameras, such that using two cameras requires twice as much time spent labeling frames, especially when each camera view observes the animal and enclosure from a different perspective. To generate a training set of 2D labeled frames, the user had to manually label frames per animal from each camera view independently. This means that each additional camera added to a rig requires that the user generate an additional set of labeled frames for that view. A keypoint can be triangulated in 3D from as few as two views (Figure 3a, 1 and 2), and LUC3D lets the remaining views inherit the reprojection of 3D position as filled-in keypoints rather than the user labeling it manually (Figure 3a, 3). The user then corrects any reprojections that do not match the animal in the video (Figure 3a, 4). Therefore, when using reprojection-aided labeling, adding additional cameras to an experimental preparation can drastically save the user time when labeling frames, while increasing the precision of each keypoint’s placement.

**Fig. 3:**
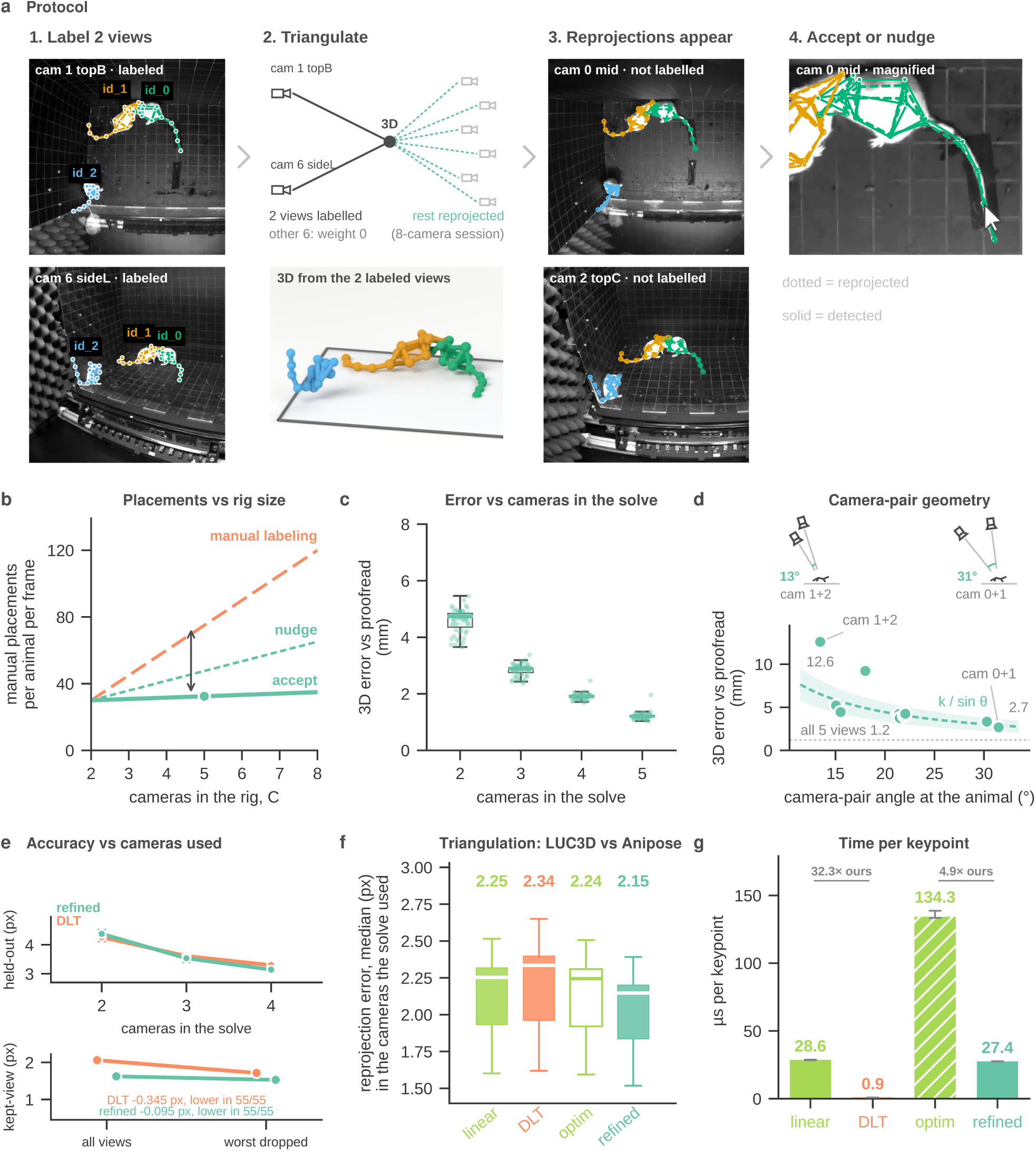
Reprojection from two labeled views reduces manual labeling, and LUC3D triangulates accurately and faster than existing tools. **a**, Reprojection-aided labeling on one frame of an eight-camera recording. 1) the user labels the keypoints in two views. 2) LUC3D triangulates each keypoint from these two views. 3) reprojects it into the other views. 4) The user accepts each reprojection or moves it onto the animal. **b**, Keypoints placed by hand per animal as cameras are added to the rig, when every keypoint is labeled by hand and when reprojections are used. Reprojections are accepted within 10 pixels (solid) or corrected to within 5 pixels (dashed) of the keypoint’s true position. The lines are a model, and the filled marker is the measured five-camera rig. **c**, Distance of the 3D keypoints from the dataset’s reference reconstruction as cameras are added to the solve. Each dot is the median of one session. The five-camera value is the distance between a single-frame solve and the reference, which is smoothed over time. **d**, Two view 3D error against the angle between the two cameras, measured at the animal. Each marker is one camera pair. The dashed curve falls as 1*/* sin *θ*, with a band of *±*25%, and the dotted line is the error when all five cameras are used. **e**, Top, reprojection error in a camera left out of the solve, as cameras are added, for the linear (DLT) and refined LUC3D solvers. Bottom, reprojection error in the cameras that were used, with all cameras and with each solver’s worst camera dropped. Error bars, 95% confidence intervals of the median (top) and the mean (bottom). **f**, Median reprojection error of each session, in the cameras used for the solve, for two LUC3D and two Anipose solvers. The LUC3D refined solver has the lowest error in all 55 sessions. Box plots show the median and interquartile range, with whiskers at 1.5 times the interquartile range. **g**, Time to triangulate one keypoint with each solver, measured on the same 200,010 keypoints for every solver. Bars are medians over timed passes and whiskers the interquartile range. Panel a shows one frame of the HardFight recording. Panels b to g use MouseDyad-10M.

Reprojection more than halves the number of keypoints placed by hand on a five-camera rig (Figure 3b). On the five-camera rig, accepting reprojections with 10 pixels of reprojection error reduces manual placements from 75 per animal to 32, a 2.3x reduction in manual labeling. Modifying those reprojections would result in 48 placements per animal, 37% fewer than manual labeling, and would reduce the reprojection error to ∼5 pixels. The saved labor grows with rig size because the labeled two views can reproject to any number of remaining cameras. This estimates the labeling effort saved by using reprojections in the labeling process. This assumes that each node would be manipulated separately, but the GUI provides methods for translating the points for one animal in a view (hereafter referred to as an “instance”) which can also reduce labeling time.

Every additional camera improves the 3D estimate. A 3D point solved from two views is a median 4.74 mm from the reference, and the full five-view solve is 1.21 mm from it (Figure 3c). The reference is the dataset’s own 3D reconstruction, which was triangulated from the same five views and smoothed over time. The value at five cameras therefore measures how far a single-frame solve sits from this smoothed reconstruction, and the values at fewer cameras are relative to it.

The cost of solving from only two views depends on where those two cameras are placed relative to each other (Figure 3d). Error decreases roughly as *k* over the sine of the angle between the two cameras relative to the animal (dashed curve). For example, the narrowest angle pair at 13 degrees gives 12.6 mm of error while the widest at 31 degrees gives 2.7 mm. The floor when all five views contribute is 1.2 mm. Strategic camera placement when setting up the rig and careful choice of the two labeled views for reprojection can also reduce labor cost. Increasing the number of contributing views leads to more accurate 3D estimates. Scored relative to a camera the solve never saw, error in the DLT solve falls from 4.26 pixels with two cameras to 3.30 pixels with four cameras (Figure 3e, top). Figure 3e shows that dropping an unreliable view can reduce reprojection error (bottom).

LUC3D’s triangulation matches the accuracy of existing tools at several times the speed. Across the solvers for LUC3D and Anipose, the median reprojection error over sessions ranges from 2.15 to 2.34 pixels (Figure 3f). Neither library’s non-linear refinement shows a large improvement for the MouseDyad-10M dataset. LUC3D’s linear and refined solvers are 32 and 4.9 times faster per keypoint than the matching Anipose solvers (Figure 3g). Including lens undistortion, which was timed separately for both libraries, the linear path is 11 times faster. The linear solver takes 0.89 *µ*s per keypoint, which supports interactive use in the browser.

### 2.4 Tracking difficulty reflects level of occlusion

SLAP-2M illustrates the difficulty of 3D tracking, from a single mouse in an empty arena to groups of mice among multiple enrichment objects. The SLAP-2M dataset was purposely collected to provide multiple challenges for 2D and 3D pose estimation systems. Each of the 74 SLAP-2M sessions varies by number of animals and number of environmental enrichment objects which increase difficulty due to occlusions (see Figure 4a for a visualization and Supplementary Video 3 for every recorded combination). A keypoint needs at least two views to be triangulated, so a keypoint that can only be placed in one or no cameras cannot be recovered by triangulation. Figure 4b shows how often occlusion leaves a keypoint impossible to triangulate. In the easiest sessions from difficulty level 1, 0.21% of keypoints cannot be recovered. The number of unrecoverable points stays below 3.3% through rating 5, then rises to 11.5% at rating 6 and 15.2% at rating 7, where 6.3% of keypoints are seen by no camera and 8.9% by only one. Every other keypoint is labeled in at least two views, so it can be triangulated and reprojected into the other views that miss it. Labeled keypoints also become less consistent as difficulty level increases (Figure 4c). The mean reprojection error of the proofread 2D labels rises 1.41×, from 3.56 pixels at rating 1 to 5.02 pixels at rating 7, and the 98th percentile doubles from 11.2 to 22.5 pixels. Figure 4d shows how the LUC3D tracker’s cross-view IDF1 degrades with difficulty. It ranges from near-perfect tracking with two animals and no enrichment objects (rating 2, median IDF1 = 0.998) to multiple animals among multiple enrichment objects (rating 7, median IDF1 = 0.836). Every stratum from difficulty level 2 through 5 has a median of 0.98 or above, which suggests near-perfect identity tracking.

**Fig. 4:**
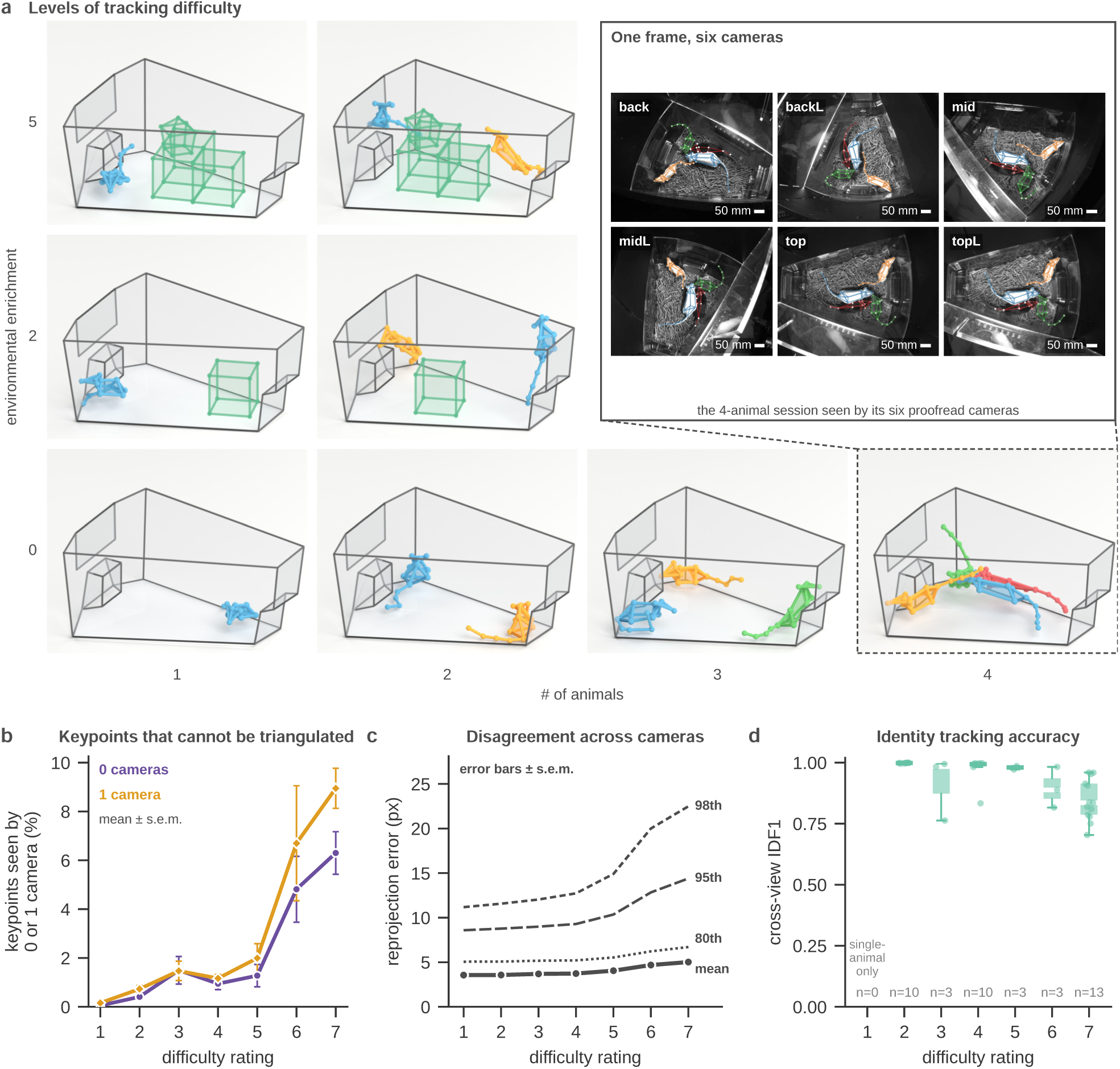
Tracking difficulty in SLAP-2M reflects environmental enrichment and the amount of occlusion. **a**, SLAP-2M sessions arranged by the number of animals and the amount of environmental enrichment, which together set each session’s difficulty rating. Each cage is one session, drawn from its own cage corners and tracked 3D poses, with the front wall shown as an outline so that the interior is visible. The inset shows one instant of the four-animal session in its six proofread cameras (scale bars, 50 mm). **b**, Percentage of keypoints that cannot be triangulated, because proofreaders could label them in no camera (purple) or in only one camera (orange) by difficulty rating. **c**, Reprojection error of the proofread 2D keypoints by difficulty rating. The solid line is the mean, the dotted line is the 80th percentile, and the dashed lines are the 95th and 98th percentiles. **d**, Cross-view IDF1 of the LUC3D tracker by difficulty rating, one point per multi-animal session. Rating 1 contains only single-animal sessions. Means in b and c are shown *±* s.e.m. across sessions. Box plots show the median and interquartile range, with whiskers at 1.5 times the interquartile range. Panels b and c use the proofread keypoints of all 74 sessions, tail keypoints excluded. B counts *∼*30.9 million 3D keypoints, and C scores their *∼*161.9 million 2D labels across the cameras. Panel d uses the 42 multi-animal sessions, tracked from SLEAP predictions as in Figure 2 (Section 4.7).

### 2.5 LUC3D reveals a geometric social motif that 2D tracking cannot quantify

Cross-view identity and metric 3D distance reveal a social behavior that 2D tracking cannot measure. We describe a social rearing behavior in the MouseDyad-10M dataset, in which two mice rear together face to face and then sniff each other while still on their hind legs (See Section 4.11 for Methods). The behavior shows a leader and follower structure in the timing of joint rearing onset. Figure 5a shows the 3D coordinates of one display beside the same instant in each of the five camera views (see also Supplementary Video 2). Every camera is mounted between 58 and 76 degrees above the animals, so no single 2D view reliably recovers the vertical axis, and single views give pixel estimates without metric distances.

**Fig. 5:**
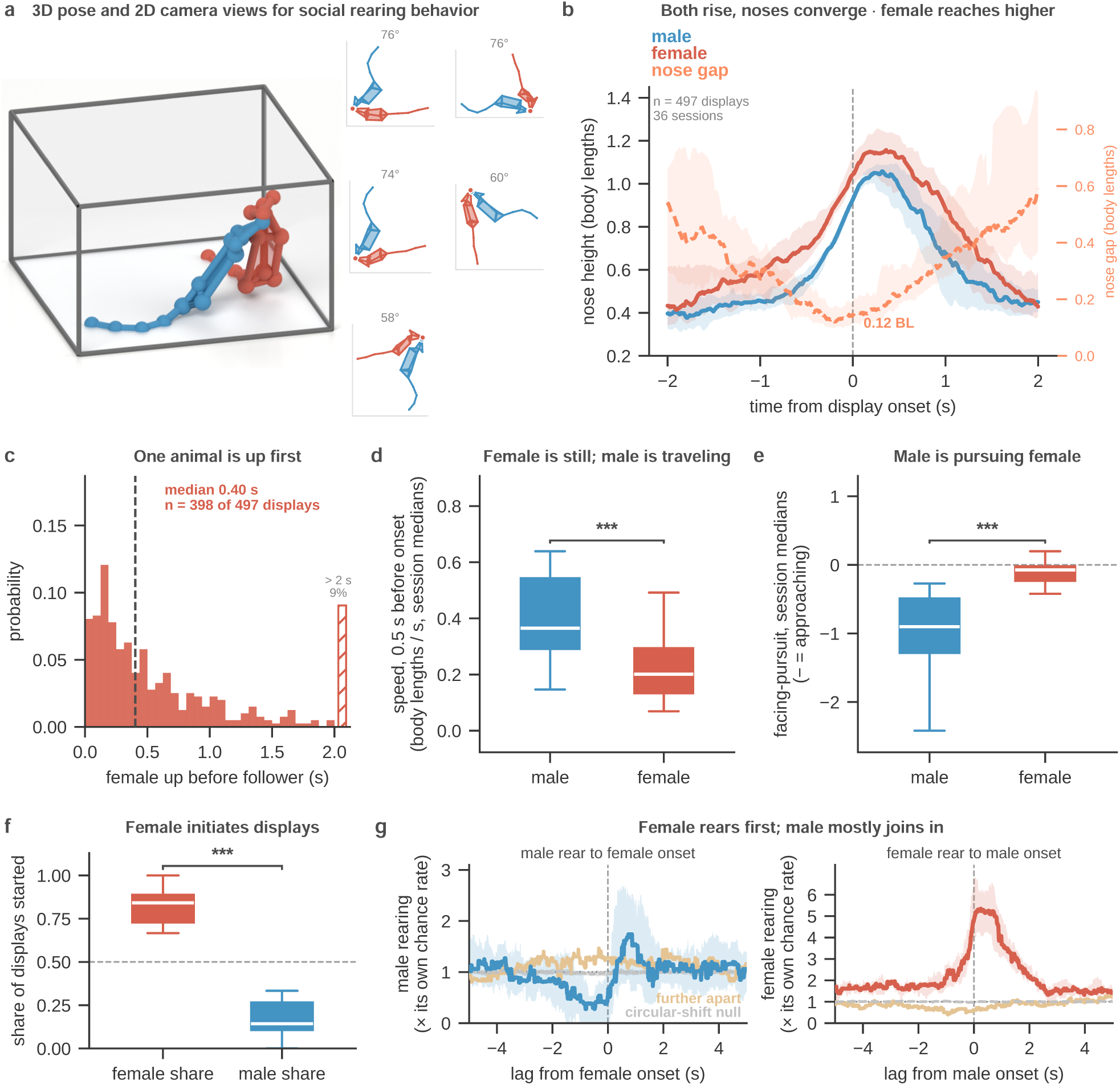
Male-female pairs rear together face to face, and the female usually leads. **a**, One social rearing display, shown as the 3D reconstruction projected into each of the five cameras. All cameras look down on the arena from 58 to 76 degrees, so the animals’ height can only be measured in 3D. **b**, Nose height of the male (blue) and the female (red), and the distance between their noses, around the start of a display. Lines are the median across sessions and bands the interquartile range. **c**, Delay between the female’s and the male’s rear onset in the 398 displays led by the female. The striped bar collects the 9% of delays longer than 2 s. **d**, Speed of each animal in the 0.5 s before a display, in body lengths per second. The male is faster (paired Wilcoxon test, *P* = 6.3 *×* 10^−4^). **e**, Pursuit score before a display, which is negative when an animal faces its partner and moves toward it and positive when it moves away. The male’s score is lower in all 36 sessions (paired Wilcoxon test, *P* = 2.9 *×* 10^−11^). **f**, Share of displays started by each sex in the 22 sessions with at least six displays (paired Wilcoxon test, *P* = 4.8 *×* 10^−7^). The dashed line marks equal shares. **g**, Probability that one animal is rearing around the other’s rear onset, relative to its own chance rate. Left, the male around female onsets. Right, the female around male onsets. The main lines are onsets with the animals within two body lengths, yellow lines onsets with the animals further apart, and gray lines a shufÒed control. The box plots in d to f summarize one value per session (36 sessions in d and e). They show the median and interquartile range, with whiskers at 1.5 times the interquartile range. Data are from MouseDyad-10M, nine mice (four males and five females) in 18 pairings, recorded with five cameras at 150 fps. A display is counted when both animals rear, with the neck above 0.75 of the body length, and their tail bases stay within two body lengths for at least 0.25 s. There are 497 displays in 36 of the 55 sessions. Panel g uses 9,073 rear onsets from all 55 sessions, 2,688 of them with the animals within two body lengths. Its curves are medians over the sessions with at least 20 onsets, 31 sessions for the male and 12 for the female.

There are 497 social rearing displays in 36 of the 55 sessions, with a median duration of 0.71 s. Both animals rise together and their noses converge just before onset (Figure 5b). In the displays led by females, the female rears a median 0.40 s before the male joins (Figure 5c). In the half second immediately before a display, the male is still traveling and the female is largely stationary (Figure 5d). Using one summary value per session, the session medians are 0.37 body lengths per second for the male and 0.20 for the female, and the male is faster in 27 of 36 sessions (paired Wilcoxon *P* = 6.3 × 10*^−^*^4^). The two animals also differ in the direction of that motion. The pursuit score is negative when an animal is oriented toward its partner and closing in (Figure 5e). Over the 1.5 s before onset the male’s session median is negative (−0.90), indicating orientation toward the female and approach, and the female’s is close to zero (−0.07), indicating neither approach nor retreat. The male’s score is more negative in all 36 sessions (paired Wilcoxon *P* = 2.9 × 10*^−^*^11^). Both differences also hold with the pair as the unit of replication. The male is faster in 13 of 14 pairs and has the more negative pursuit score in all of them (Section 4.11). Across individual animals, each of the four males scores more negative than each of the five females (Mann-Whitney *P* = 0.016). With only nine animals, the speed difference across animals is not significant (Mann-Whitney *P* = 0.19).

The sequential ordering of the displays is asymmetric, and in this cohort the female is the leader. The female initiates 398 of the 493 displays (80.7%) and starts more displays than the male in 35 of the 36 sessions with at least one display. The male initiates the remaining 95 displays. Four displays in which both animals rose on the same frame have no initiator. The one exception is an exact tie, and no session has the male as the outright leader. Over the 22 sessions with at least six displays, the female’s median share of displays is 0.84 (paired Wilcoxon *P* = 4.8 × 10*^−^*^7^, Figure 5f).

The 55 MouseDyad-10M sessions are repeated recordings of nine mice (four males and five females) in 18 pairings, and the displays are spread unevenly over them, from one to 305 per female, so the analysis was repeated with the pair as the unit of replication. Every one of the 17 distinct pairings that produced at least one display has the female as its leading member. Over the 14 pairs with at least five displays, covering 494 displays, the female starts a median 0.81 of that pair’s displays and leads in all 14 of 14 (sign test *P* = 1.2 × 10*^−^*^4^). Pooled over these pairs, the female starts 395 of the 490 displays with an initiator (80.6%).

Control tests rule out simple explanations for the female leader (Supplementary Section 5.1). The female’s lead is not explained by a higher rate of rearing. Across the 22 sessions with at least six displays, the female’s proportion of the pair’s rearing time does not predict how often the female starts a display (*r* = −0.022, *P* = 0.92). The result also does not depend on how rearing is detected. An animal counts as rearing when its neck is higher than 0.75 of its own body length, so the threshold scales with each animal’s size. An absolute 60 mm threshold applied to both animals returns the female as leader in all 22 sessions. Body size does not explain the effect either. The male is usually the longer animal, but in the eight sessions where the female is the longer animal, the female still starts 78% of displays (binomial *P* = 6 × 10*^−^*^9^).

We found that rearing is sequentially coupled when the animals are in close proximity. The median likelihood of rearing relative to the other animal’s rearing onset demonstrates an asymmetric leader and follower relationship (Figure 5g). The male’s rearing probability is below baseline prior to and during the female rear onset and reaches 1.7 times the baseline rate ∼0.8 s later. The probability of the female’s rearing rises prior to the male onset, reaches 4.8 times the baseline rate when the male starts, and peaks at 5.3 times a quarter of a second later. When the male starts a rear within two body lengths, the female is already rearing 53.3% of the time. When the female starts one, the male is already rearing 5.9% of the time. The male therefore usually joins a rear the female has already started. This sequential relationship is much weaker when the animals are further apart.

## 3 Discussion

LUC3D makes 3D multi-animal pose estimation accessible with no installation required. It speeds up the two most manual steps of a 3D project, reprojection-aided labeling and proofreading. Reprojection-aided labeling cuts manual placements by more than half on a five-camera rig, and the cross-view tracker resolves identity across every camera at once, so a session is proofread once rather than once per camera. We provide two large open source datasets, with 65 million video frames, of multi-animal multi-view 3D pose estimation for examining social behavior: MouseDyad-10M and SLAP-2M. Finally, we have also shown that LUC3D can reveal a social behavior that can only be quantified in 3D.

To resolve the cross-view identity tracking problem, previous work suggested that exhaustive hypothesis testing would be required. Unfortunately, this method becomes increasingly untenable with each additional animal or camera view [18]. However, an alternate method is greedy Hungarian matching with a 3D correspondence term. We found that on equivalent frames, greedy Hungarian matching misgroups fewer frames than the exhaustive hypothesis testing strategy when pooled over all sessions, and runs over an order of magnitude faster. The ablation shows that the 3D term is what makes the greedy search competitive, since without it the greedy search falls behind exhaustive enumeration. At 0.28 to 0.69 ms per frame, it is fast enough for interactive cross-view tracking in a browser-based GUI.

In order for LUC3D’s pipeline to incorporate each step of 3D pose estimation, a fast, built-in triangulation function was required. Compared to Anipose, LUC3D’s linear triangulation runs at 32 times the speed (0.89 *µ*s per keypoint, excluding undistortion), fast enough for interactive use in the browser and performs with comparable accuracy. We also incorporated our new calibration tool, calibrat3, which achieves a lower median reprojection error than Anipose on all rigs tested.

LUC3D uses labeling redundancy to reduce labor and minimize manual annotation and provides a method for accepting or modifying reprojections from a small number of labeled camera views to the larger number of remaining views. Reprojections can themselves serve as labels, and because they derive from a single triangulated point, they are 3D-consistent across all views. This is most useful for occluded keypoints, which reprojection fills in. Reprojection-aided labeling allows the generation of training sets from scratch using far fewer manual labels than a traditional single-view set up.

For multi-animal datasets, correcting 2D identity switches during proofreading is one of the largest remaining blocks of manual labor. Without LUC3D it scales linearly with the number of cameras (a two-camera rig doubles the proofreading time and a four-camera rig quadruples it, since each additional video would traditionally need to be separately proofread). Identity must also be associated across views for the same animal to be recognized as the same ID in every camera, which takes additional proofreading. The LUC3D cross-view tracker instead produces an automated proofread output for both cross-view identity assignment and temporal identity tracking. As a result, the user needs only to review the session once, reducing proofreading labor. Using more cameras gives the LUC3D cross-view tracker more redundant evidence, so it makes fewer errors that require manual correction. On MouseDyad-10M its within-view IDF1 rises from 0.92 with two cameras to 0.97 with four or five (Section 4.8.1). The GUI also provides visual aids and tools which make the errors easy to catch and correct. The tracker recovered identity in rats as well as mice, from geometry alone and without using the markings on the animals. That test used reprojected 3D poses, so its performance on a real detector’s output in another species remains to be measured.

We also demonstrated how 3D pose estimation can be used to reveal leader-follower dynamics during male-female interactions in mice. This analysis shows that close proximity rearing has a social structure [22–24]. It also revealed an asymmetry between the two animals of each pair, the female leading and the male following, that would not be visible without cross-view identity and metric distance. The female consistently leads the display, starting the large majority of them across nearly every session and every pairing. The male’s approach in the moments beforehand looks like directed pursuit. The male stays oriented toward and closing in on the female while the female is largely stationary. Whether this asymmetry is a sex difference or a property of these nine animals cannot be settled with four males and five females, and whether it is related to courtship would require independent measures such as ultrasonic vocalizations, estrous staging, or mating assays. The result establishes a stable directional social dynamic in this cohort, and multi-camera 3D pose estimation was necessary to characterize it.

LUC3D has limitations that suggest future directions. For instance, it requires calibrated and synchronized cameras. It also relies on 2D detections from a model trained and run on a GPU, although every later step runs in the browser. This still lets one shared GPU serve many lab members, who can label and proofread from their own laptops. Tracking accuracy falls in the most enriched SLAP-2M sessions, where objects and other animals hide keypoints from several cameras at once (median cross-view IDF1 0.84 at rating 7, where 15% of keypoints are seen by fewer than two cameras). Very large sessions can also become impractical due to the limited memory available to a browser tab.

Quantifying how behavior unfolds in 3D will be critical for the future of behavioral and computational neuroscience. Accurate 3D poses enable downstream applications such as embodied biomechanical modeling [25, 26]. These body models learn to reproduce lab behaviors in physics simulators, and they need large amounts of 3D pose data to train. By reducing manual labor and eliminating installation requirements, LUC3D facilitates the next generation of neuroscientists to understand behavior in its natural 3D form.

## 4 Online Methods

### 4.1 Browser-based 3D annotation and tracking GUI

LUC3D is a multi-view pose annotation and 3D proofreading interface that runs entirely in a web browser (Supplementary Video 4 walks through the full workflow). The GUI is written in JavaScript served as static files with no installation and no build step, either from the hosted deployment or from any local static file server. A project is composed of a set of local files that the user opens, saves, and loads.

The application loads multi-camera video with frame-accurate WebCodecs decoding, synchronized scrubbing across all views, and camera calibrations in TOML or JSON format. SLEAP predictions are read lazily, so prediction files larger than the browser’s available memory can still be opened. Several recording sessions can be held open at once, each in its own dockable pane. A project can also be opened with no video at all, as a skeleton together with imported 3D points, in which case the whole duration of the reconstruction remains navigable.

The arena floor can be placed in 3D by labeling its corners in two views and triangulating them with the Define Planes feature, which also sets the origin of the coordinate frame (Supplementary Video 4). Skeleton instances and their keypoints are created and edited directly on the video frames. Skeleton nodes and edge definitions can be edited inline from its own panel. Nodes can be added, renamed, or removed, and edges rewired. A project has one skeleton, shared by all its sessions. Reprojection-aided labeling protocol (quantified in Figure 3) can be performed with the interface. The user labels two views, the 3D point is solved from those two alone, and the reprojection appears in every remaining view to be accepted or nudged. Figure 3b counts *C* × *N* placements for manual labeling and 2*N* plus the corrected reprojections for reprojection-aided labeling. This assumes that each node would be manipulated separately, but the GUI provides methods for translating the points for one animal in a view (hereafter referred to as an “instance”) which can also reduce labeling time.

A cross-view tracker groups the detections that belong to the same individual across views and assigns one identity per animal per frame (see Figure 2 and Supplementary Figure 4). A guided dialog exposes the tracker’s per-node weights, per-view inclusion and cost thresholds (Section 4.8). This allows the tracker parameters to be chosen for each user’s particular dataset. Tracks and identities are created per session so they can be matched across camera views.

Triangulation can be performed as a linear solve or a non-linear refinement (Section 4.9), with optional rejection of any keypoint observation whose reprojection error exceeds a threshold. The reconstruction is shown in an interactive Three.js viewport that also shows the position and field of view of each calibrated camera. The viewport can be placed at any camera’s viewpoint for direct comparison with that camera’s video, and can be exported as an MP4. The proofreader’s working signal is the reprojection error. Every view shows its own detections together with the current 3D reconstruction reprojected into that view, drawn as a dotted skeleton with different node markers so the two are not confused. The reprojection errors are also listed in an Instance info panel on the right side of the screen which appear as green, yellow and red depending on the size of the error (See Supplementary Figure 3 for GUI screenshot). A timeline widget in the style of SLEAP contains frames, tracks and labeled regions for navigation, and the keyboard shortcuts are listed and can be rebound in the settings dialog.

Three instance types are distinguished throughout the interface, namely user instances, predicted instances, and reprojected instances. User instances are created or edited directly by the annotator and appear as solid fully editable skeletons. Predicted instances are produced by the cross-view tracker or imported from SLEAP which are read-only until converted into user instances. Reprojected instances are computed by projecting the current 3D reconstruction back into each 2D view and are also read-only until converted into user instances. After a new user instance is created, its keypoints are dragged into position individually from scratch or moved together as a rigid group to reposition an already roughly correct skeleton based on the previous frame for that view. Each node can also be cycled through visible or occluded with a right-click. Instances can be deleted individually or users can use the custom instance delete prompt to systematically delete instances by specifying the particular label type, view, session and/or frame.

The information panel reports the reprojection error for every instance group in each contributing camera. A reprojected instance in an unlabeled view can be converted to an editable instance with a double-click and adjusted in place. The group can then be re-triangulated to improve correspondence between labels and reprojections. The reprojection overlay is the proofreader’s principal diagnostic to identify consistent labels across views. An annotation that diverges visibly from its dotted reprojection indicates a mislabeled keypoint, a node swap, an identity swap, or a calibration error.

The 3D viewport can also be exported as an MP4 video. An export dialog makes output resolutions available from 480p to 2160p and reports the frame count and estimated file size before rendering begins. The renderer encodes the 2D overlays and 3D triangulated points with WebCodecs directly in the browser.

Projects are written as SLEAP .slp files. The files include the columnar session data introduced in format version 2.8, which contain the calibration, the cross-view identities and the 3D points. This ensures that a project can save and load round-trip through SLEAP and through this application without data loss. Labels can also be exported as JSON, 3D points as HDF5, and the 3D viewport as MP4.

### 4.2 Software

The application is written as ES modules in JavaScript with no build step. It reads and writes version 2.8 of the SLP format through sleap-io.js and HDF5 through h5wasm, and renders in three dimensions with Three.js. A recent Chromium-based browser is recommended, because frame-accurate decoding uses WebCodecs. The benchmark drivers run the unmodified application modules under Node 26 for batch comparison. Analysis and figures were produced in Python 3.12 with NumPy, pandas, h5py, SciPy and Matplotlib. Tracking metrics come from motmetrics, comparison triangulation from aniposelib 0.7.2 and comparison calibration from aniposelib 0.8.0 [7, 27, 28]. The speeds quoted in the Introduction, up to 16,030 frames per second for triangulation and 14,694 for tracking, were measured in the application itself (v1.0.5) in Chrome on a 16 GB Apple M1 MacBook Pro, on a MouseDyad-10M session, with a timing snippet run in the browser console. The benchmarks in Figures 2g and 3g were run on a different machine with a different method, so their numbers differ from these.

### 4.3 Datasets

Table 1 describes the details of the two large multi-camera datasets that are made available in this paper: MouseDyad-10M and SLAP-2M. The SLAP-2M dataset was created with eight CD-1 mice (Charles River Strain Code 022), four C57BL/6J mice (Jackson Laboratory Strain Number 000664), and four BALB/c mice (Taconic Biosciences BALB/cAnNTac). The MouseDyad-10M dataset used all CD-1 mice. SLAP-2M was manually proofread in 3D. For MouseDyad-10M, identities were proofread in each camera (see below). In addition, the HardFight dataset was used to test social interaction with a high incidence of occlusions. Its session has three CD-1 mice from a cohort of 12, mostly littermates. In general, the experiments consisted of placing mice into an enclosure for 1 to 60 minutes and observing behavior. All mice were kept on a reverse 12/12 dark/light cycle.

**Table 1:**
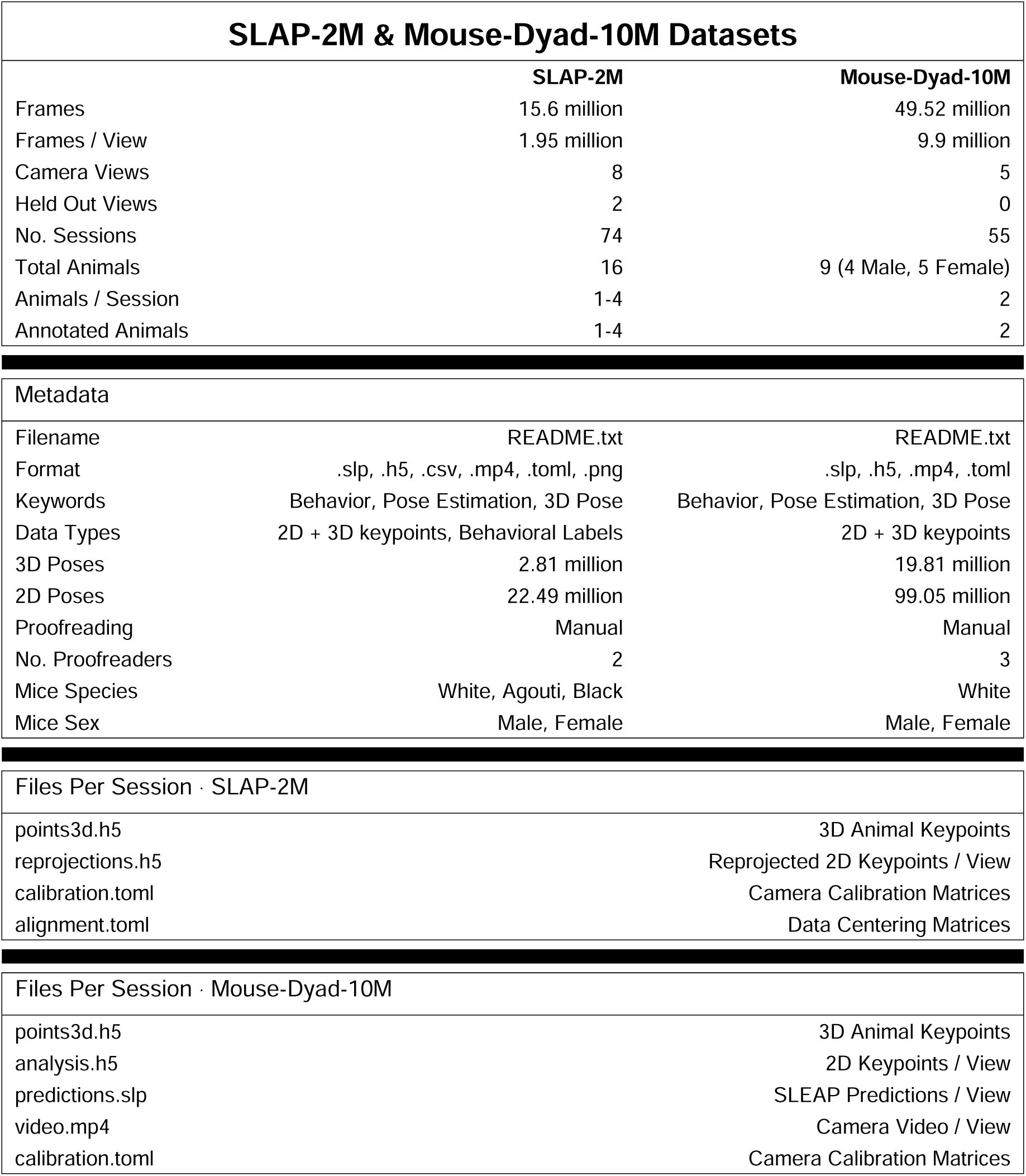
Dataset datasheets. Composition, metadata and per-session file inventory of the two datasets: SLAP-2M and MouseDyad-10M.

|  | **SLAP-2M** | **Mouse-Dyad-10M** |
|---|---|---|
| Frames | 15.6 million | 49.52 million |
| Frames / View | 1.95 million | 9.9 million |
| Camera Views | 8 | 5 |
| Held Out Views | 2 | 0 |
| No. Sessions | 74 | 55 |
| Total Animals | 16 | 9 (4 Male, 5 Female) |
| Animals / Session | 1-4 | 2 |
| Annotated Animals | 1-4 | 2 |
| Metadata |  |  |
| Filename | README.txt | README.txt |
| Format | .slp, .h5, .csv, .mp4, .toml, .png | .slp, .h5, .mp4, .toml |
| Keywords | Behavior, Pose Estimation, 3D Pose | Behavior, Pose Estimation, 3D Pose |
| Data Types | 2D + 3D keypoints, Behavioral Labels | 2D + 3D keypoints |
| 3D Poses | 2.81 million | 19.81 million |
| 2D Poses | 22.49 million | 99.05 million |
| Proofreading | Manual | Manual |
| No. Proofreaders | 2 | 3 |
| Mice Species | White, Agouti, Black | White |
| Mice Sex | Male, Female | Male, Female |
| Files Per Session · SLAP-2M |  |  |
| points3d.h5 | 3D Animal Keypoints |  |
| reprojections.h5 | Reprojected 2D Keypoints / View |  |
| calibration.toml | Camera Calibration Matrices |  |
| alignment.toml | Data Centering Matrices |  |
| Files Per Session · Mouse-Dyad-10M |  |  |
| points3d.h5 | 3D Animal Keypoints |  |
| analysis.h5 | 2D Keypoints / View |  |
| predictions.slp | SLEAP Predictions / View |  |
| video.mp4 | Camera Video / View |  |
| calibration.toml | Camera Calibration Matrices |  |

The MouseDyad-10M corpus consists of 55 sessions, each recording two mice with five synchronized cameras at 150 frames per second for approximately 20 minutes. Each session contains a male and female mouse (N = 9, 4 male, 5 female) in 18 unique pairings. The males always occupy track 0 and the females always occupy track 1. A 15-node mouse skeleton was used throughout, comprising the nose, the left and right ears, the head, neck and trunk, the left and right shoulders and haunches, the tail base, and four tail nodes, three along the tail and one at its tip. The five cameras are elevated above the arena and identified by serial number. The corpus contains ∼10 million synchronized frames per view, ∼20 million 3D instances and ∼100 million 2D instances reprojected into the five views. Identities were proofread in each camera before triangulation. Brief identity swaps between cameras remain in 0.2% of frames. Figures 2f to i, 3b to g and 5 and Supplementary Figure 4a were measured on this corpus.

The SLAP-2M data repository contains ∼2.8 million 3D poses and ∼22.5 million 2D poses reprojected from them into the eight views. The multi-animal dataset was collected during 74 recording sessions with 8 camera views with between 1-4 animals in each session. The eight cameras (back, mid, side, top, back-left, mid-left, side-left and top-left) were pointed at the center of the arena. Side and side-left were left out of the analysis because of high incidence of occlusion that reduced the 2D pose estimation quality. However, the side views are included as a challenge for future methodologies. Both males and females are included in the dataset as well as multiple mouse strains (white, black, and agouti). This dataset is unique in its level of diversity and emphasis on freely roaming behavior. The dataset includes six levels of enrichment, namely none, interactive planar, single animal with one vertical object, multiple animals with one vertical object, multiple animals with several vertical objects, and multiple animals with several vertical and opaque objects. While many popular 2D datasets lack environmental enrichment, adding levels of enrichment highlights the need for 3D pose estimation. By adding more aspects of the environment which the animals can climb, 3D pose becomes more relevant. This makes the dataset an ideal candidate for benchmarking new models for behavioral clustering (both supervised and unsupervised), 3D pose estimation, and ReID tracking. Each arena has either black or white bedding. Each session also has an assigned difficulty rating from 1 to 7 which increases with the number of animals and enrichment objects in the arena. The difficulty rating is the stratifying axis throughout Figure 4. Figures 2a and b, Figure 4, Supplementary Figure 4b to d and the 42 multi-animal sessions of Figures 2f and g were taken from SLAP-2M.

The third recording, HardFight, is a single eight-camera, three-mouse session at 60 frames per second, from which a 300-frame window was taken for Figure 1a and Figure 3a. Figure 2e is one frame from LUC3D’s reconstruction of all 36,000 frames of the session.

To test the cross-view tracker on a second species, we also used three published rat datasets from social-DANNCE [13]. TRIADS, BEDDING and SCN2A together contain 41 sessions of two or three rats recorded by six calibrated cameras at 50 frames per second. The social-DANNCE animals are marked with food coloring to keep track of their identities, and these marks were not used here. The procedure is described in Supplementary Section 5.5 and the result is in Supplementary Figure 4e.

### 4.4 Animal Ethics Statement

All animal procedures were approved by the Princeton University Institutional Animal Care and Use Committee and the Salk Institutional Animal Care and Use Committee and were in accordance with National Institutes of Health Standards.

### 4.5 Video acquisition

Videos were acquired with FLIR Blackfly S cameras at 30 to 150 fps, using either SpinView (Spinnaker SDK) or campy (https://github.com/ksseverson57/campy) on a Windows 11 computer. The video resolution for all three pose datasets is 1280×1024 pixels. The SLAP-2M video data has a frame rate of 30 FPS, except the 13 sessions which are 60 FPS. The MouseDyad-10M data has a framerate of 150 FPS. The HardFight dataset has a framerate of 60 FPS.

We also provide Panopticon, a Python acquisition app for hardware-triggered multi-camera recording with Basler cameras. Panopticon previews every camera and records trigger-synchronized video with real-time GPU encoding. It monitors capture lag during recording so that dropped or delayed frames are flagged while the session is still running. Panopticon also has an on-board interface for optogenetic stimulation paradigms, using the same trigger board as acquisition for exact correspondence between output capture signals and output stimulation. Synchronization was tested with an infrared LED driven from the stimulation output and visible to all nine cameras (Supplementary Section 5.9). Each session is written to a LUC3D compatible structured folder with metadata describing the animals, rig and acquisition machine. Panopticon also captures and solves camera calibrations with live coverage feedback, as described in Section 4.6.

### 4.6 Camera calibration

Calibration is the process by which camera parameters and 3D points are iteratively estimated by comparing reprojected 2D points with multi-view ground truth. The ground truth is estimated by using an easily detected stimulus, such as a ChArUco board with a checkerboard pattern and bit encodings called ArUco patterns. A calibration is performed by pointing multiple cameras at a central point, and the ChArUco board is presented in multiple orientations and positions visible from all cameras. Then sparse bundle adjustment is used to estimate optimal camera parameters and 3D points which are associated with unique positions of the detected markers. The camera calibration is stored in Anipose-compatible TOML format which records the camera name, resolution, intrinsics, extrinsics (rotation and translation), and distortion parameters. Reprojection errors are calculated between the 3D triangulated checkerboard patterns and the 2D detections and the residual is measured for each iteration throughout optimization. All 3D coordinates are expressed in millimeters. Calibrations for the SLAP-2M and MouseDyad-10M dataset were performed in Anipose. This paper presents a browser-based calibration tool called calibrat3 which takes an OpenCV/Anipose-inspired calibration suite and makes it more accessible without any installation required (https://talmolab.github.io/calibrat3/). The tool also allows for backwards compatibility with deprecated versions of OpenCV’s ChArUco boards and the newest formats. Calibrat3 refines the camera intrinsics, extrinsics and board poses together with sparse Levenberg–Marquardt bundle adjustment. It is written in JavaScript with multiple options for camera distortions and performs per-camera intrinsics calculations. Its accuracy was benchmarked against Anipose on three different camera rigs. The benchmark methods can be found in Supplementary Section 5.6. The results can be found in Supplementary Section 5.8 and Supplementary Figure 2.

Panopticon (Section 4.5) also performs calibration directly from the acquisition window in realtime (Supplementary Figure 1). A ChArUco board is carried through the arena while a live coverage display shows which cameras and camera pairs have detected the board and which quadrants of each view remain uncovered, and it signals when every camera has sufficient shared detections and spatial coverage. This replaces the guesswork of deciding when enough footage has been collected, since poorly constrained intrinsics otherwise appear acceptable and fail only near the frame edges. Once the calibration is recorded, Panopticon solves for the intrinsics and extrinsics of every camera, writes the result in the same Anipose-compatible TOML format described above, and copies it into the session folder so that each recording has its own calibration. The at-rig solve confirms the recording and the camera geometry before animals are placed in the arena, and the final calibration should be solved from the saved video by calibrat3 or Anipose. To measure the accuracy of the at-rig solve, a ChArUco calibration recorded through Panopticon on a nine-camera Basler rig (Supplementary Figure 1) was solved by Panopticon and by calibrat3. Both calibrations were scored on the same 758,665 corner observations seen by two or more cameras, using the reprojection scoring of Supplementary Section 5.6. The results are reported in Supplementary Section 5.9.

### 4.7 Two-dimensional detection pool

Within each benchmark every method received the same per-frame detections, with all track and identity information removed, so the comparisons isolate cross-view association and triangulation from the quality of the underlying detector. SLEAP, whose tracker is part of its own pipeline, was scored on its own tracking of the same predictions. Ground-truth identities were transferred onto the pool by matching detections to the proofread instances frame by frame and camera by camera at an intersection-over-union threshold of 0.5. With all identity labels stripped before the methods ran, the comparison reflects each method’s own association quality given the detector’s spatial accuracy. SLEAP produced the detections (Supplementary Section 5.7). LUC3D reads SLEAP .slp files and neither trains nor runs a detector of its own. In both datasets the pool is SLEAP’s predictions before proofreading with none of sleap-nn’s instance filters applied, minus every detection with fewer than five visible body keypoints. The four tail nodes do not count toward the five. Identities in both datasets are scored without them, on the ground truth and every tracker alike. On MouseDyad-10M every tracker used the tail nodes for tracking, including in ByteTrack’s bounding boxes. On SLAP-2M, SLEAP and ByteTrack did not use them, and ByteTrack’s boxes were built from the body nodes only. LUC3D gave them zero weight in its association costs (Section 4.8).

Behavioral analyses and tracking metrics (Figures 2h and i, 4d and 5 and Supplementary Figure 4) were computed on every frame of each session, with Figures 2h and i scored on the frames without missing detections. Figures 2f and g use the eligible frames and samples described in Section 4.8. The per-keypoint geometric measurements in Figures 4b and c were also computed on every frame. Figures 3b to g were computed on every 15th frame across the same 55 sessions as the tracking benchmarks. LUC3D’s values come from the application’s own solvers. Subsampling was required because these measurements are per-keypoint and combinatorial in the number of cameras.

The camera-subset comparisons in Figure 3c to e require combinatorial solves across camera subsets, so they utilized the subsampled data. At 150 fps an animal typically moves less than a millimeter between consecutive frames, so sampling every 15th frame (0.1 s intervals) reduces redundant information and increases computational tractability. The stride does affect temporal smoothing, which is why that term is disabled in the triangulation comparison.

### 4.8 Cross-view association

Figure 2a shows a diagram of the 3D and 2D correspondence terms and their retained per-view anchor nodes. For each frame the application maintains a set of 3D targets and solves one Hungarian assignment per camera against the same frame-start targets, re-triangulating once all cameras are assigned, at a cost of *O*(*C* · *A*^3^) for *A* animals and *C* cameras (Figure 2c). The cost of pairing a target *t* with a detection *d* is summed over the skeleton nodes *k* with per-node weights *ω_k_*, and then negated to allow for the solver to minimize it. On SLAP-2M the four tail nodes were given zero weight, so the association costs rely on the 11 body nodes. The tail nodes still enter the epipolar matching that starts a new track, and the triangulated poses. On MouseDyad-10M every node kept a weight of 1. In the app every node has a weight of 1 by default. The two-dimensional term compares the detection with the target’s projection *π*(*t*) into that camera, in undistorted ideal-pinhole coordinates,

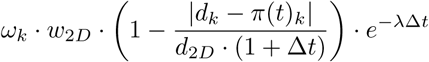

where *w*_2*D*_ is the weight of the 2D term, *d*_2*D*_ its distance limit, Δ*t* the target’s age and *λ* the time penalty, so age both widens the distance allowance and decays the whole term. The three-dimensional term is on the perpendicular distance from the target’s 3D point to the ray back-projected from the detection, with no age decay,

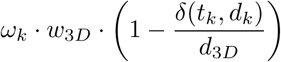

Here *δ*(*t_k_, d_k_*) is the distance from the target’s 3D point to the line of sight through the detection, *w*_3*D*_ is the weight of the 3D term and *d*_3*D*_ its distance limit. The ratio *r* = *w*_3*D*_*/w*_2*D*_ sets how strongly the association relies on 3D geometry relative to 2D image distance. The app and Figure 2h label the two weights corr3d and corr2d. A retained per-view detection older than *N* frames is evicted from a target’s 3D anchor before that frame is matched. The age of a retained detection is called staleness and *N* is the eviction window. An anchor is therefore only ever triangulated from cameras that saw the animal recently, so views that have lost the animal no longer pull the anchor toward stale positions. Supplementary Section 5.3 reports the parameter search over *N*. By default *w*_2*D*_ = 1, *w*_3*D*_ = 6, *d*_2*D*_ = 10 in normalized image units, *d*_3*D*_ = 25 mm, *λ* = 0.1 per frame and *N* = 20 frames. The same defaults were applied without change to the social-DANNCE rat datasets (Supplementary Section 5.5).

We compared the greedy Hungarian method with a re-implementation of the exhaustive procedure of Maree et al. [18]. It enumerates all (*A*!)*^C^* groupings of a frame’s detections into identities, triangulates and reprojects each one with LUC3D’s own triangulation code, and keeps the grouping with the lowest error. The published procedure cannot carry an identity from one frame to the next. To score its identity switches and IDF1, each frame’s groups were matched to the identities of the previous enumerated frame with the Hungarian algorithm on the distance between their 3D centroids. This step does not change which detections are grouped together, so Figure 2f does not depend on it. Like LUC3D’s association costs, it uses the four tail nodes on MouseDyad-10M and leaves them out on SLAP-2M. Exhaustive enumeration needs exactly *A* detections in every camera, so only frames meeting that condition were scored. Of the ∼10.6 million frames in the 97 multi-animal sessions (55 MouseDyad-10M and 42 SLAP-2M), ∼9.9 million were eligible. To keep enumeration tractable, the three-animal SLAP-2M sessions were enumerated on five of their cameras (back, back-left, mid, mid-left and top, 7,776 hypotheses per frame) and the four-animal sessions on three (back, mid and top, 13,824 hypotheses per frame). The greedy method was run on the same cameras. The two-animal configurations took 4.0 to 5.4 ms per frame, and every eligible frame was enumerated. The three- and four-animal configurations took 0.87 and 1.5 s per frame, so 2,000 and 1,000 eligible frames per session were sampled uniformly across the whole session rather than from its start.

For Figure 2g both methods were timed on the same frames, drawn at random from the eligible frames, up to 1,000 per session with two animals and 100 with three or four. The two methods ran one at a time and in alternation, over three rounds, and the median over the rounds is reported for each method. Only the solve was timed, headless in Node.js 26 on the modules of LUC3D v1.0.5, not in a browser. Each timed run starts a new process, so LUC3D’s time includes its start-up compilation. No other job ran during the timing. The processor time of every process was measured every 5 s, and the calls during which other processes used more than half a core were timed again. The host’s one-minute load average, from work outside this study, had a median of 6 on its 64 cores. Four animals in six cameras need ∼190 million hypotheses per frame, far above our cap of 10^6^, so exhaustive enumeration was not run there. The greedy time was measured the same way on 100 eligible frames per session. The exhaustive time in Figure 2g (open marker) is estimated from the fastest rate per hypothesis measured in any configuration, so the true time would be longer. The bar on that marker spans two estimates. Its low end treats a relabeling of all the animals at once as the same grouping and uses the fastest rate. Its high end counts every relabeling and uses the rate of the largest configuration that was run.

Grouping quality in Figure 2f compares which detections are grouped together, not the identity labels assigned to each group, so a consistent relabeling is not counted as an error. Both methods were scored on the same ∼9.9 million frames that were enumerated, with the four tail nodes excluded from the ground-truth matching. Ground-truth identities were matched to each detection at an intersection-over-union (IOU) threshold of 0.5. Exhaustive enumeration needs exactly one detection per animal in every camera, so all of these frames meet that condition.

The sweep in Figures 2h and i fixes *w*_2*D*_ = 1 and varies *w*_3*D*_ to sample twenty values of *r* from 0 to 24 over all 55 MouseDyad-10M sessions at full length. Switches are reported per 100,000 camera frames for the detections used for each comparison. Both methods are scored on the 47.9 million camera frames without missing instances, out of 49.5 million in the full sessions.

#### 4.8.1 Camera-subset identity

To measure how many cameras identity tracking needs, the tracker was re-run on subsets of *k* = 2, 3 and 4 of the five MouseDyad-10M cameras, over the same 55 sessions and the same detections. For each *k* we used three fixed subsets of cameras, the same for every session, so that differences between sessions are not mixed with differences between subsets. Running every possible subset on every session was not feasible. The full five-camera run reproduces the main 55-session result exactly. Identity tracking holds up with fewer cameras. Mean within-view IDF1 is 0.92 with two cameras, 0.96 with three, 0.97 with four and 0.97 with all five. Over the 55 sessions, the three two-camera subsets give 158 to 250 identity switches, the three-camera subsets 86 to 298, the four-camera subsets 92 to 128, and the full rig 130. Fewer cameras means fewer camera frames, so switches are compared as rates per camera frame.

### 4.9 Triangulation

The application’s default linear solver undistorts the observations into ideal pinhole camera coordinates and minimizes the algebraic error in closed form by direct linear transformation. The non-linear solver minimizes a geometric error in each camera’s native, still-distorted pixels while holding the cameras fixed, using a soft-L1 loss followed by an L1 polish phase and initialized from the linear solution, and it never accepts a step that increases the error. The application labels the non-linear solver “Refined (Ref)”.

The application can optionally drop, for each keypoint, observations whose reprojection error is above a threshold and re-solve that keypoint from the remaining views. The bottom of Figure 3e does not use this threshold. As an analysis protocol it drops the single worst-fitting view of every keypoint, re-solves the keypoint from the remaining views with the application’s solvers and scores the all-view solve and the re-solve on the same remaining views. This shows a small but consistent decrease in reprojection error. The error at the top of Figure 3e is measured in a camera that was left out of the solve.

The comparison triangulation method in Figures 3f and 3g is aniposelib 0.7.2, which corrects for lens distortion with OpenCV and then solves each point on the CPU. GPU-based alternatives were not included because of the browser-based constraint. The four solvers were compared in pairs matched by linear and non-linear algorithm classes. The library’s optional outlier-rejection mode was more than 70 times slower than its linear solver on the same keypoints, measured in earlier sittings, and was not included. The temporal smoothing term in the non-linear path was disabled. The smoothing penalty assumes consecutive entries are consecutive video frames, but the input is sampled every 15th frame and filtered to keypoints visible in all five views. Consecutive entries are therefore separated by at least 0.1 s of real motion, and the penalty treats that motion as noise. With smoothing enabled, the library’s non-linear solver has a higher held-out error than its own linear solve in all 55 sessions.

Solve times exclude undistortion for every solver because one library undistorts inside the triangulation call and the other outside it. The excluded undistortion is about 1.7 microseconds per keypoint for this work. For aniposelib it is about 0.4 microseconds, measured in an earlier sitting on the same keypoints. All solvers were timed in one sitting on a single thread pinned to one core, alternating between the two libraries over five rounds, with no other process running. The two fastest steps take about 0.2 to 0.3 s per pass, so they were timed over 20 consecutive passes in each round. Each bar is the median of its timed passes and the whisker is the interquartile range. LUC3D’s solvers ran headless in Node.js 26 on the modules of LUC3D v1.0.5, not in a browser. The two libraries are written in different languages and run on different runtimes, so this comparison is of two implementations as a user would encounter them.

Figure 3c reports the 3D distance to the reference reconstruction in millimeters. The reference was not made with LUC3D. A per-frame triangulation of the same detections, made independently of LUC3D, lands a median 1.25 mm from it, close to LUC3D’s 1.21 mm, and the reference changes a median 23 times less from frame to frame, so it was smoothed over time. These values are over the 39 sessions in which that independent triangulation is usable. The reference also has its own noise (median reprojection error about 2.4 pixels), but this enters every camera count equally. These values show relative improvement with camera count, not absolute 3D accuracy. Figure 3d repeats the two-camera solve for each camera pair and plots its error against the angle between the two cameras. The triangulation analyses in Figure 3 use all 15 nodes, including the tail. In one session (20250904_140306), one camera’s SLEAP predictions fragment into 12 short tracks after frame 34,447. The tracker input is identity-free and unaffected, but the analyses in Figure 3b to g read the first two tracks of each camera, so they use only frames 0 to 34,447 of that session.

### 4.10 Occlusion and reprojection error

Figure 4b measures how many keypoints occlusion leaves impossible to triangulate. A keypoint is one node of one animal in a single frame. The proofread 3D pose is complete in every frame of all 74 SLAP-2M sessions, so a keypoint is never missing in 3D, but it can be missing from a view when the proofreaders’ 2D label in that camera does not carry it. Each proofread 2D instance in a camera is paired with a proofread 3D animal reprojected into that camera by Hungarian matching on mean keypoint distance, and for each keypoint we count how many of the six analyzed cameras carry it. A keypoint carried by no camera or by exactly one cannot be triangulated by any subset of these cameras. Every keypoint carried by two or more can be triangulated from them and reprojected into the views that miss it. The four tail nodes are excluded, which leaves 11 body nodes and ∼30.9 million keypoints over the 74 sessions. Each session contributes one percentage, and each difficulty rating is summarized as the mean ± s.e.m. between its sessions (*n* = 4 to 13 sessions per rating). Figure 4c uses the same matching and keypoints. Its reprojection error is the pixel distance between each labeled 2D keypoint and the proofread 3D keypoint reprojected into that camera. Each session’s mean error is averaged within a rating, with the s.e.m. between sessions. Each percentile line (80th, 95th and 98th) is that percentile of each session’s errors, averaged within a rating.

### 4.11 Behavioral analysis of social rearing

Rearing is defined as an increase in head height above the floor while on the two hind legs, often with the forelimbs supporting the animal’s weight against a wall. We were particularly interested in the social case, when rearing occurs as a synchronized pair interaction. This behavior begins as a synchronous rear in close proximity followed by orofacial sniffing while the animals are still on their hind legs. We refer to one such event as a social rearing display.

A frame was scored as reared when the animal’s neck height exceeded 0.75 of its own body length, where body length is the median nose-to-tail-base distance for that animal over the session. Rearing bouts shorter than 0.25 s were discarded, and gaps of 0.15 s or less within a bout were merged. A social rearing display required both animals to be rearing with their tail bases within two body lengths of each other for at least 0.25 s. Nose height and nose gap in Figure 5b are expressed in body lengths. A display begins only when the second animal rears up, and the leader is the animal whose bout began earlier. A display in which both bouts begin on the same frame has no leader and is excluded from the initiation counts, which applies to four of the 497 displays. The lag in Figure 5c is the interval between the two bout onsets, computed over female-led displays only.

Speed was calculated for each animal in body lengths per second, taken as the median over the 0.5 s before display onset (Figure 5d). We computed a direction vector by subtracting the partner’s position from the animal’s position, and a pursuit score as the dot product of the animal’s body orientation vector with that direction vector scaled by the animal’s own speed (Figure 5e). Each display’s pursuit score is the median over the 1.5 s before onset, divided by the animal’s median speed over the session. A negative score means the animal is oriented toward its partner and approaching it, and a positive score means it is oriented away and retreating. For Figures 5d and e the unit of replication is the session. Each animal’s values were reduced to one median per session over that session’s displays, and the two animals were compared with a paired Wilcoxon signed-rank test across the 36 sessions with at least one display. The comparison was repeated with the pair as the unit over the 14 pairs with at least five displays, and across animals, with each mouse’s median over every display it took part in and the four males compared with the five females by a Mann-Whitney U test. The displays are unevenly spread over animals, from one to 305 per female and 16 to 212 per male, which is why a test over displays would overstate the evidence.

Figure 5f shows each animal’s share of initiations, one value per session, and a session’s leader is the animal that initiated more displays. The female’s and the male’s shares were compared with a paired Wilcoxon signed-rank test across sessions. Pooled shares were computed over displays, weighting each session by its display count. Sessions with fewer than six displays were excluded from the share comparison in Figure 5f. The rearing-rate and height-threshold controls used the same sessions. The pair-level analyses used a threshold of five displays, which keeps 14 of the 17 pairs.

The 55 MouseDyad-10M sessions are repeated recordings of 9 mice in 18 pairings, so the analysis was also run with the pair as the unit of replication. Every session of a pair was aggregated into one observation, and pairs with fewer than five displays were excluded, leaving 14 pairs covering 494 displays. Pairs were tested with a sign test on which animal led and a Wilcoxon signed-rank test of the female’s share using a null value of 0.5.

The rearing-rate and height-threshold controls were run over the 22 sessions with at least six displays, the same sessions as in Figure 5f. For the rearing base rate, each session’s fraction of female-initiated displays was correlated with the female’s fraction of the pair’s total rearing time. For the height threshold, the detection was re-run with an absolute 60 mm neck height applied to both animals in each pair. It was also re-run with a single threshold shared by both animals of a pair, set at 0.75 times the pair’s mean body length. To control for body size, each animal’s median body length over the session was used to identify the longer animal of the pair. Male and female body length were compared with a paired Wilcoxon test across the 36 sessions with at least one display. Whether the leader was the shorter animal was tested with a sign test across the 35 sessions with a unique leader. Sex is confounded with the track slot in this corpus, since the male occupies slot 0 and the female slot 1 in every session. To check that the detector has no slot preference, every session was re-run with the two slots exchanged in the input. Swapping the slots gave identical results, with each animal keeping its own values.

For Figure 5g we took every rearing onset by one animal and computed the probability that the other animal was rearing at each lag within 5 s of that onset, divided by the second animal’s base rate of rearing. Onsets were split by the tail-base separation between the animals at onset into those within two body lengths and those further apart. A null distribution was generated by circularly shifting the second animal’s rearing time series, 24 shifts per ordered pair. The shift preserves that animal’s rearing rate, bout durations and autocorrelation, and removes only the temporal alignment between the two animals. Sessions contributing fewer than 20 onsets in a condition were excluded from that condition.

### 4.12 Tracking metrics

IDF1 measures how accurately a tracker maintains the correct identity of each animal over time [21]. It is computed as the number of detections matched to the right individual divided by the average of the ground-truth and predicted detection counts, which weights false positives and false negatives equally. Identity switches occur when the ID for one instance is mistaken for another relative to ground truth labels.

IDF1 and identity switches were computed with the motmetrics library on the shared detection pool [27, 28]. Within-view IDF1 was computed separately for each camera and averaged. Cross-view IDF1 pools all cameras into a single accumulator with one global identity per animal. LUC3D asserts one identity per animal across cameras and its ids are pooled directly. SLEAP and ByteTrack track each camera on its own and number their tracks from zero in every camera. A “track 0” in one camera therefore has no relation to a “track 0” in another. We keep their track ids separate per camera. Which animal becomes track 0 in a camera reflects detector confidence and not any cross-view reasoning. A tracker whose ids are kept separate per camera can match the truth in at most one camera, which puts it near 1*/C*. Error terms in Supplementary Figure 4d are reported as percentages of camera frames. False negatives were measured but are not plotted, since they dominate every method’s error budget. Figure 4d reports the cross-view IDF1 of the LUC3D tracker on the 42 multi-animal SLAP-2M sessions. Detections are matched to ground truth by the bounding boxes of their keypoints. For every analysis, these boxes leave out the four tail nodes.

### 4.13 Baseline configuration for tracking comparison

The tracking comparison is reported in Supplementary Section 5.2 and Supplementary Figure 4a to d. For Supplementary Figure 4a to d, SLEAP and ByteTrack were run with settings that improve their performance over the defaults. LUC3D is given the *n* global identities the user specifies, so it cannot create more. SLEAP by default maintains an unbounded track pool, and ByteTrack, as run in the benchmark, retires any track lost for more than 60 frames, which is 1 to 2 s for SLAP-2M and 0.4 s for MouseDyad-10M. Both original settings fragment identities on long sessions.

SLEAP was limited to as many tracks as there are animals in the session, two for MouseDyad-10M and one to four for SLAP-2M. It ran in sleap-nn 0.3.0 for MouseDyad-10M and sleap-nn 0.2.0 for SLAP-2M. SLEAP tracked the same detections as the other two trackers (Section 4.7). It applied no detection filters of its own in either dataset.

ByteTrack’s default of ending a track after 60 unmatched frames often ends tracks too early, so this was turned off. Its tracks were then merged into one identity per animal. Each new track takes an identity that is not already in use in that frame. If more than one is free, it takes the identity whose last bounding box overlaps its first box the most, with ties broken by the distance between box centers. The track keeps that identity until it ends, so ByteTrack’s own frame to frame matching is unchanged. The same changes were used for SLAP-2M, with tracks merged into the session’s number of animals. After the merge a frame can hold two rows with the same identity, and these duplicate rows are scored as they are. This raises ByteTrack’s IDF1 slightly above 1 in 10 of the 444 SLAP-2M camera-sessions (at most 1.0049), which is an artifact of the scorer rather than better tracking.

### 4.14 Anchor staleness and the fresh-anchor tracker

The cross-view tracker associates identities between cameras using 3D correspondence [10]. For each identity and each camera, the most recent matched detection is retained for comparison. The tracker re-triangulates a 3D anchor for each identity from all retained detections. Chen et al. [10] weight each retained detection by an exponential decay in age (their Eq. 11), but the reference implementation computes the weights without applying them, and computes the sign of the exponent such that older detections would receive larger weights if the routine were completed. All retained detections therefore enter the triangulation at equal weight regardless of age.

Modifications were made to the anchor update to reduce identity switches. Staleness eviction deletes every retained detection older than *N* frames at the start of each frame, so the anchor is triangulated only from cameras that saw the animal recently. *N* was swept over 1, 10, 20 and 30 frames and compared with no eviction. The sweep is shown in Supplementary Figure 4f and reported in Supplementary Section 5.3. A target with fewer than two retained detections keeps its last anchor without re-triangulating. Under synchronous scoring, every anchor is frozen at the start of a frame. All views are scored from the frozen anchors, and retriangulation happens once at frame end. This prevents a wrong assignment in one view from contaminating the scores of the remaining views. Also, the distance limit of the 3D term was reduced from 50 to 25 mm. The anchor update was otherwise unchanged from the reference implementation.

The sweep in Supplementary Figure 4f was run on all 74 SLAP-2M sessions with identical per-camera detections, and the same eviction windows were also run on the 55 MouseDyad-10M sessions. Paired comparisons across sessions used the two-sided Wilcoxon signed-rank test. The configuration selected by these runs is the current default.

### 4.15 Statistics

The session is the unit of analysis throughout unless stated otherwise. For MouseDyad-10M, the 55 sessions are repeated recordings of nine mice (four males and five females) in 18 pairings. For SLAP-2M the individual animals are not identified in the metadata, so no finer unit is available. Error distributions are summarized by their median and interquartile range. The exceptions are Figures 4b and c, which report the mean ± s.e.m. across the sessions of each difficulty rating. Bands from the 25th to the 75th percentile show the spread across sessions of each session’s median. The comparisons in Supplementary Figure 4a report 95% confidence intervals over sessions. The per-session comparisons of direction in Supplementary Figure 4c use sign tests. The paired comparisons in Figures 5d to f use one value per session and a Wilcoxon signed-rank test across sessions, with pair-level and animal-level repeats in Section 4.11. The temporal coupling in Figure 5g is compared with circular-shift nulls. No correction for multiple comparisons is applied across figures.

### 4.16 3D renders and animations

The 3D renders in Figures 1a and b, 2a, b and e, 3a, 4a and 5a, and in Supplementary Videos 1 to 3, were made in Blender 5.2 through its Python module (bpy). Every scene is built by a script from the data, with no manual placement, so each render can be regenerated exactly. The renders in Figures 1b, 2a and b, 4a and 5a show each dataset’s own 3D poses as shipped, and those in Figures 1a, 2e and 3a show LUC3D’s reconstruction. Scenes were rendered with the Cycles path tracer on an NVIDIA A40 GPU, with denoising. Frames were encoded with ffmpeg as H.264 video at 1920 × 1080 and 30 frames per second.

### 4.17 AI Declaration

The code for LUC3D, the software tool presented in this paper, and for calibrat3, the browser-based calibration tool (both written in JavaScript), and the scripts used to analyze the data and generate figures (written in Python) were developed iteratively with Claude Opus (Anthropic) acting as a coding assistant. The authors designed the software, the experiments, the comparisons and the analyses, and specified them in detail. Claude implemented them in code under the authors’ direction. The authors reviewed and tested all of this code and verified that it performs the intended analyses and that the outputs accurately represent the underlying data. Claude also served as an editing assistant for the manuscript. The scientific questions, interpretation of results and conclusions are the authors’ own. The authors take full responsibility for the content of this manuscript.

## Data Availability

The pose estimation data from the SLAP-2M dataset is available at https://huggingface.co/datasets/talmolab/slap-2m. The pose estimation data from the MouseDyad-10M dataset is available at https://huggingface.co/datasets/talmolab/mouse-dyad-10M. The social-DANNCE rat data are available at https://dataverse.harvard.edu/dataverse/socialDANNCE_data [13].

## Code Availability

The LUC3D GUI can be accessed directly at https://luc3d.sleap.ai/. The code for LUC3D can be found at https://github.com/talmolab/luc3d with documentation at https://talmolab.github.io/luc3d-docs/.

The calibrat3 browser-based calibration GUI can be accessed directly at https://talmolab.github.io/calibrat3/. The code for calibrat3 can be found at https://github.com/talmolab/calibrat3 with documentation at https://talmolab.github.io/luc3d-docs/#camera-calibration.

The Panopticon acquisition and live calibration app can be accessed at https://github.com/talmolab/panopticon.

The scripts, plot-ready tables and measurement deposits that generate every figure in this paper are available at https://github.com/talmolab/figures-luc3d.

## Acknowledgments

We thank Elly Lewis, Ivan C. Alcantara, Kee Wui Wang, Akira Nagamori, Ayesha Thanawalla, Oren Wilcox, Max Wang, and Eiman Azim for helpful discussions and software testing. This work was supported by a L.I.F.E. Foundation grant to T.D.P., a Salk Innovation Award to T.D.P., an NIH BRAIN Initiative grant RF1MH132653 to T.D.P., an NIH BBQS grant to T.D.P., A.L.F. and M.M. (U01DA063581), and an R37MH102441 (NIMH) to K.M.T., who is an HHMI Investigator and the Wylie Vale chair at the Salk Institute for Biological Studies. Additional funding was from DP2MH126375 (A.L.F.), NIH R01MH126035 (A.L.F.), the New York Stem Cell Foundation (A.L.F.), the Simons Collaboration on the Global Brain (A.L.F.), the Klingenstein Foundation (A.L.F.), an NINDS R35 Research Program Award R35NS111580 (M.M.), the Princeton Neuroscience Institute (M.M.), an EMBO Postdoctoral Fellowship ALTF 1084-2021 (B.M.) and a Princeton Neuroscience Institute C.V. Starr Fellowship (B.M.). A.L.F. is a New York Stem Cell Foundation Robertson Investigator.

## Author Contributions

E.J.L., S.N.O., A.L.F. and T.D.P. conceived the project. T.D.P. and A.L.F. supervised the project, M.M. and K.M.T. supervised the work carried out in their laboratories. T.D.P. developed LUC3D prototype. L.M. laid the algorithmic groundwork for the LUC3D backend. E.J.L. designed LUC3D and calibrat3 and implemented calibrat3. E.J.L., T.D.P., S.N.O., and J.P. implemented LUC3D, and E.J.L. and T.D.P. developed its backend in sleap-io.js. I.T., A.Z. and T.W. developed Panopticon under the supervision of K.M.T., who also provided expertise on the behavioral analysis. I.T. generated Supplementary Figure 1. S.N.O. collected the SLAP-2M dataset under the supervision of A.L.F., and S.N.O., S.A. and O.T. proofread and curated it. B.M. collected and curated the MouseDyad-10M dataset under the supervision of M.M., and B.M., J.J. and E.W. proofread it. E.J.L. performed the analyses and generated the figures. T.D.P., A.L.F., M.M. and K.M.T. acquired funding. E.J.L. wrote the initial draft, and E.J.L., S.N.O., I.T., A.L.F. and T.D.P. reviewed and edited the manuscript.

---

**Supplementary Figure 1:**
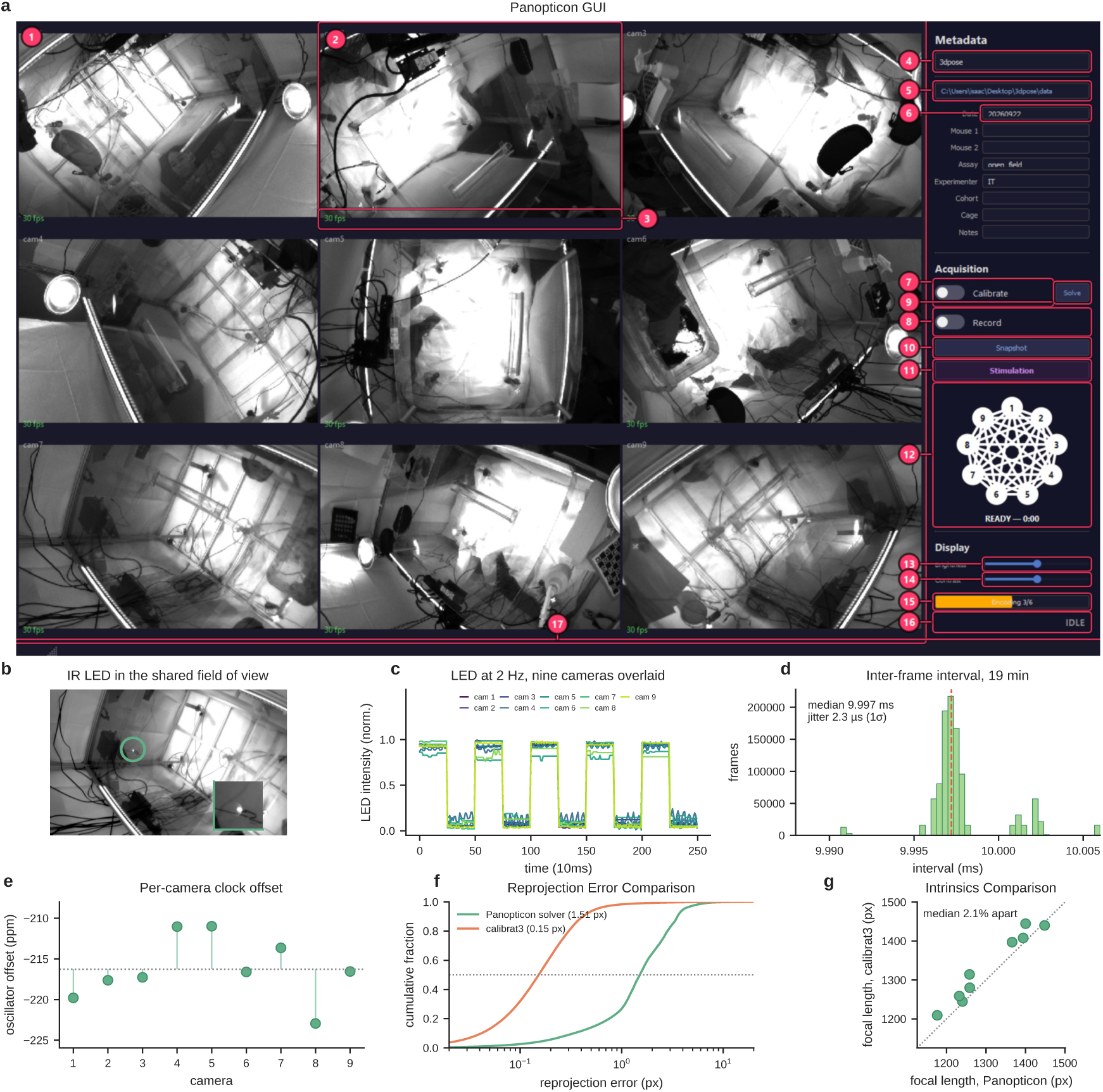
Panopticon acquires hardware synchronized multi-camera video. All panels come from one nine-camera rig of Basler a2A1920-165g5m cameras at 1920 *×* 1200, hardware-triggered by an Arduino Mega 2560. The synchronization recordings in c to e ran at 100 fps, and those in a and f at 30 fps. **a**, The acquisition window during a live nine-camera session. Left, one live preview pane per camera. Right, the sidebar with session metadata, calibrate and record toggles with the solver entry point, a snapshot button, the stimulation editor, the live calibration-coverage graph with its per-camera nodes and readiness state, and display controls above a status line. 1, camera preview grid; 2, camera pane; 3, real time frame rate; 4, profile; 5, output directory; 6, session metadata; 7, calibrate toggle; 8, record toggle; 9, calibration solve; 10, snapshot; 11, stimulation editor; 12, live calibration-coverage graph; 13, preview brightness; 14, preview contrast; 15, encode progress; 16, session state; 17, status bar. **b**, One camera view of the arena with infrared LED tied to stimulator, circled. The inset shows the lit LED. The LED is driven from the trigger board’s stimulation pin by a 2 Hz, 50% duty paradigm. **c**, LED intensity against trigger ordinal for all nine cameras over time. All nine cameras register the LED, 542 to 2,118 times above the per-pixel background at that frequency. **d**, Histogram of each camera’s inter-frame interval over a 19-minute recording. The dashed line marks the median. **e**, Each camera’s clock rate as parts per million of the nominal 100 Hz, showing the crystal spread across the array. The dotted line marks the array mean. **f**, Cumulative reprojection error for a 30 fps calibration recorded through Panopticon, solved by Panopticon’s solver vs calibrat3 on 758,665 ChArUco corner observations seen by two or more cameras. **g**, Focal length recovered by each solver, one point per camera. The dotted line marks equality. The two agree to a median of 2.1%.

**Supplementary Figure 2:**
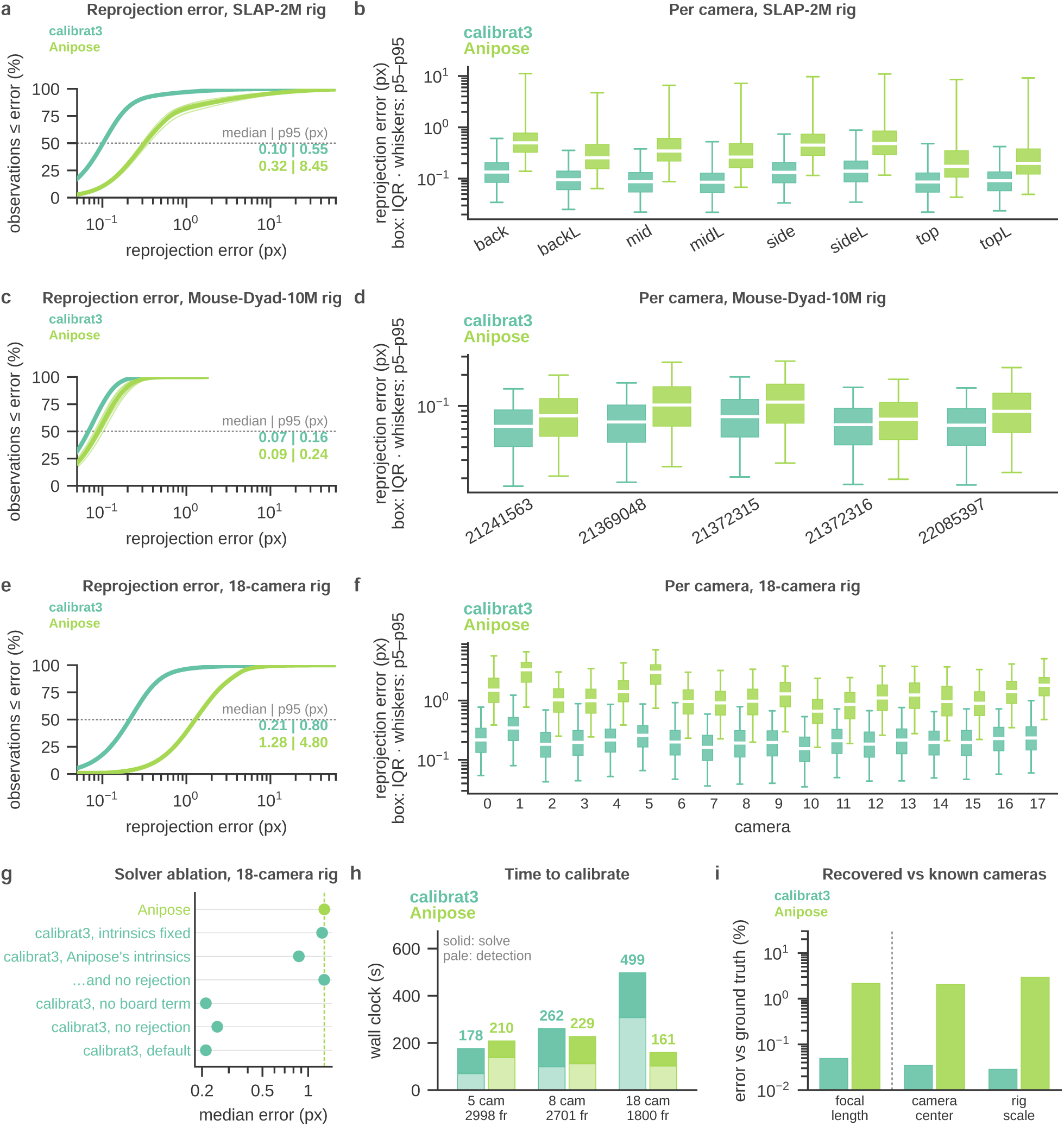
Calibration accuracy of calibrat3 and Anipose on three camera rigs. **a**, Cumulative reprojection error for calibrat3 and Anipose on the SLAP-2M rig, pooled over four calibration recordings, with each recording also drawn as a thin line. The dotted line marks the median, and the numbers give each tool’s median and 95th percentile error in pixels. **b**, Reprojection error for each of the SLAP-2M rig’s eight cameras. Boxes show the median and interquartile range, and whiskers reach the 5th and 95th percentiles, here and in d and f. **c**, Cumulative reprojection error on the MouseDyad-10M rig, pooled over five recordings. This rig was held out when choosing calibrat3’s default settings. **d**, Reprojection error for each of the MouseDyad-10M rig’s five cameras. **e**, Cumulative reprojection error on the 18-camera rig, from one recording. **f**, Reprojection error for each camera of the 18-camera rig. **g**, Median reprojection error on the 18-camera rig when calibrat3’s solver is re-run on the same detections with one setting changed at a time, such as fixing the intrinsics or using Anipose’s simpler intrinsic model. The dashed line is Anipose. **h**, Time to calibrate one session on each rig, split into board detection (pale) and solving (solid), with each tool running alone on the same machine. **i**, Percent error in focal length, camera center and rig scale on a synthetic rig where the true cameras are known. Camera center error is relative to the spacing between cameras. (Panels a to g are scored on the corners Anipose itself detected, so the comparison does not favor calibrat3. The SLAP-2M rig is 9,029 frames over four sessions at 1280 *×* 1024, the MouseDyad-10M rig 12,621 frames over five sessions at the same resolution, and the 18-camera dataset 1,800 frames at 1680 *×* 1200, all showing one 8 *×* 11 ChArUco board with 24.0 mm squares. Anipose is aniposelib 0.8.0.)

**Supplementary Figure 3:**
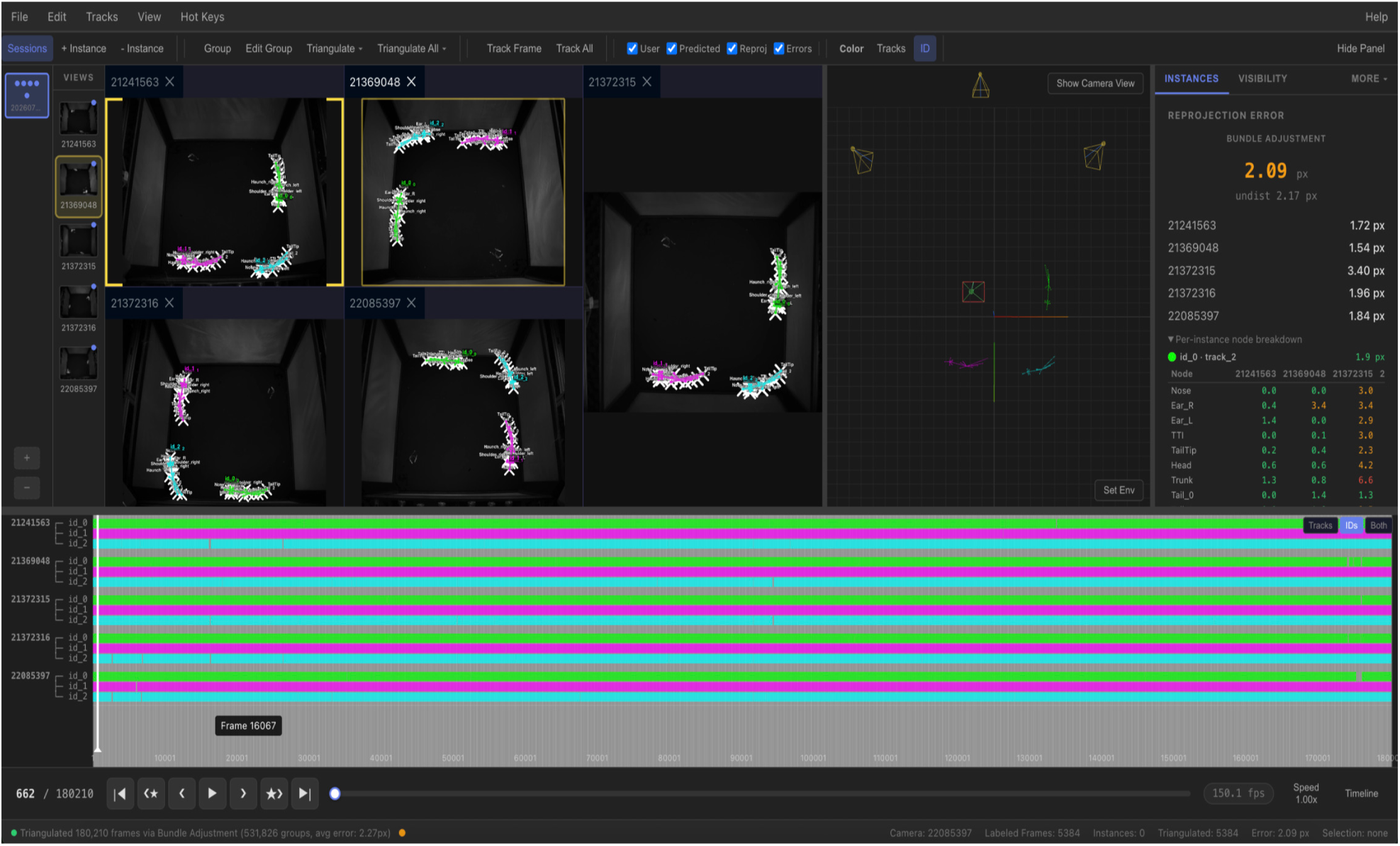
The LUC3D interface. A five-camera recording of three mice at 150 fps on the MouseDyad-10M rig, separate from the dataset, after cross-view tracking and triangulation of every frame. Top, the menu and toolbar, with controls to add and group instances, triangulate one frame or the whole session, track, toggle the user, predicted and reprojected instances and the reprojection-error overlay, and color skeletons by track or by identity. Left, the open sessions and the list of camera views. Center, the synchronized camera views in dockable panes, each showing its detections with their resolved identities and the reprojection of the 3D poses. Right of the views, the 3D viewport with the calibrated cameras and the triangulated poses. Far right, the Instances panel, which reports the reprojection error of the selected group overall, in each camera and for each node. Bottom, the timeline, with one row per identity in each camera showing the frames in which that identity is present, above the playback controls and a status bar that summarizes the triangulation.

**Supplementary Figure 4:**
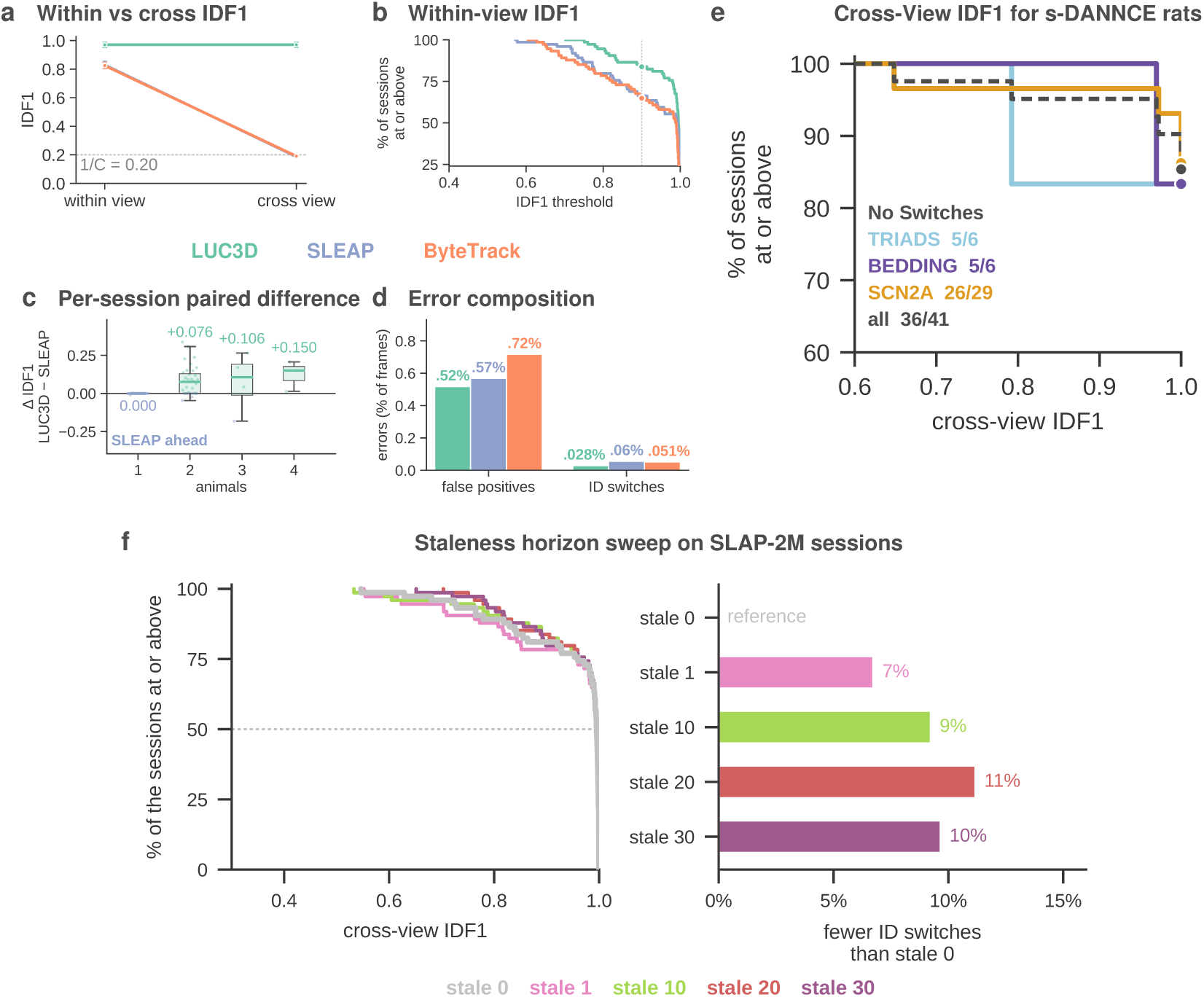
Within-view identity, cross-view identity, and 3D parameter search performance comparisons. **a**, Mean within-view and cross-view IDF1 for LUC3D, SLEAP and ByteTrack on MouseDyad-10M, with 95% confidence intervals. **b**, The share of SLAP-2M sessions whose within-view IDF1 is at or above each threshold, with one step per session. Each session’s score is the mean over its six cameras. **c**, The within-view IDF1 difference between LUC3D and SLEAP in each SLAP-2M session, grouped by the number of animals. Values above zero mean LUC3D scored higher. Boxes show the median and interquartile range, whiskers reach 1.5 times the interquartile range, and each dot is one session. **d**, False positives and identity switches on SLAP-2M, each as a percentage of camera frames. **e**, LUC3D’s performance on s-DANNCE on cross-view IDF1, one curve per dataset, with all 41 sessions as the dashed line. The counts under “No Switches” are the sessions in each dataset with no identity switch. **f**, Sweep of the staleness horizon (the anchor node eviction window) on SLAP-2M. Left, cross-view IDF1 as a survival curve, one curve for no eviction (stale 0) and for eviction windows of 1, 10, 20 and 30 frames. Right, the reduction in identity switches relative to no eviction (stale 0). Panel a uses 55 MouseDyad-10M sessions. Panels b to d use all 74 SLAP-2M sessions, *∼*11.7 million camera frames, tracked from SLEAP predictions. Panel e uses 41 social-DANNCE rat sessions with six cameras and no added noise. Panel f uses all 74 SLAP-2M sessions, *∼*11.7 million camera frames, with sessions paired across settings.

**Supplementary Videos** All four videos are available at https://talmolab.github.io/luc3d-page/.

**Supplementary Video 1. SLAP-2M.** Flythrough of a home-cage recording, showing the enclosure and its cameras rendered in 3D, four mice tracked with per-animal identity colors, and the same skeletons projected onto one camera’s footage. 13 s.

**Supplementary Video 2. Social rearing in MouseDyad-10M.** A male (blue) and a female (red) rearing together in one MouseDyad-10M session, played in real time. The top left panel shows the 3D poses, the top right panel the same poses projected onto one camera’s video, and the bottom left panel a side view against nose height. The bottom right panel plots nose height around each rear peak for the displays in this clip (thin lines) and the mean of all 497 social rearing displays in the dataset (thick lines). 14 s.

**Supplementary Video 3. SLAP-2M difficulty.** Each recorded combination of animal count (one to four) and environmental enrichment rating, one SLAP-2M session each, rendered in 3D from its own cage geometry and tracked poses. Played at 2× speed. 63 s.

**Supplementary Video 4. LUC3D walkthrough.** The full LUC3D workflow in the browser on a MouseDyad-10M session. The calibration is loaded, so the cameras appear in 3D, and the arena floor is defined from four corners and a center labeled in two views, which places the origin and grid on the floor. A skeleton is built, and an animal is labeled in two views, grouped and triangulated, with the reprojection error guiding corrections. Track All then gives each mouse one identity across all five cameras, flagged encounters are reviewed in the ID Switches tab, and an identity is reassigned from the Instances panel. Finally, the 3D poses are inspected from each camera’s viewpoint and in a free orbit. 10 min 9 s.

## 5 Supplementary Material

### 5.1 Control tests for the female-led asymmetry in social rearing

A series of control tests were performed to rule out the obvious alternatives to a female-led asymmetry in sequential behavior. One alternative is that the female leads only because of a higher incidence of rearing. Across the 22 sessions with at least six displays, the fraction of female-initiated displays was unrelated to the female’s fraction of total rearing time (*r* = −0.022, *P* = 0.92). Another alternative is that the result is an artifact of the per-animal threshold based on body length. An absolute 60 mm height threshold applied to both animals returns the female as leader in all 22 sessions. A single threshold shared by both animals of a pair returns the female as leader in 19 of 22. Finally, the finding could have been trivially explained by body size. The males are longer, with a median body length of 90.2 mm and the females 84.2 mm, and the male is longer in 28 of 36 sessions (paired Wilcoxon *P <* 0.0001), so the smaller-bodied sex is the one leading. Across the 35 sessions with a unique leader the leader is the shorter animal in 27 (sign test *P* = 0.0019), and in the eight sessions where the female is the longer animal, the female still starts 78% of displays (binomial *P* = 6 × 10*^−^*^9^). These findings indicate that the leader and follower asymmetry during social rearing is not explained by rearing frequency, threshold, or body size.

### 5.2 Cross-view identity tracking compared with per-camera trackers

Supplementary Figure 4 compares the association performance of LUC3D with the SLEAP and ByteTrack tracking methods. Supplementary Figure 4a uses 55 MouseDyad-10M sessions (five cameras, two animals). Supplementary Figure 4b to d use the SLAP-2M sessions, which range from one to four animals, and the same SLEAP and ByteTrack settings as Supplementary Figure 4a (Section 4.13). SLEAP and ByteTrack were not designed to keep identity across cameras, so the cross-view comparison demonstrates that capability and is not a benchmark of the trackers. The within-view comparison is reported to show that adding cross-view association does not cost within-view accuracy (Supplementary Figure 4a).

Within a single camera of SLAP-2M, all three methods are broadly competitive, with median within-view IDF1 of 0.995 for LUC3D, 0.992 for SLEAP and 0.987 for ByteTrack (Supplementary Figure 4b). The within-view difference between LUC3D and SLEAP is negligible in the single-animal sessions, where there is nothing to associate (median difference 0.000, mean −0.0003, Supplementary Figure 4c), although where the two differ there SLEAP is marginally higher, in 14 sessions to LUC3D’s 1. Over all 74 sessions the sign test finds no difference between the two trackers. LUC3D is higher in 32 sessions, SLEAP is higher in 24 and 18 are tied (sign test *P* = 0.35). With two or more animals LUC3D is higher in 31 of 42 sessions, SLEAP in 10, and one is tied (sign test *P* = 0.0015), with a median difference of 0.076 at two animals. Only seven sessions have three or four animals, where cross-view association should help most. LUC3D is higher in six of seven (sign test *P* = 0.13), by a median of 0.11 with three animals and 0.15 with four. Most errors in all three trackers are missed detections, 80 to 84% of each tracker’s errors (Supplementary Figure 4d). Of the rest, LUC3D has the fewest false positives (0.52% of camera frames, against 0.57% for SLEAP and 0.72% for ByteTrack) and about half the identity switches (0.028%, against 0.056% and 0.051%).

The difference appears when identities are pooled across cameras (Supplementary Figure 4a). Scoring all five MouseDyad-10M cameras into one accumulator with a single global identity per animal, LUC3D’s IDF1 is unchanged, 0.971 within view and 0.971 across. The per-camera trackers retain about 23% of their within-view IDF1 (SLEAP 0.829 to 0.186, ByteTrack 0.825 to 0.190). A tracker that labels each camera independently can match the ground truth in at most one camera, so its cross-view IDF1 cannot exceed about 1*/C*. The comparison therefore scores SLEAP and ByteTrack on a capability they never claimed. It shows that the capability is present in LUC3D and absent in per-camera trackers.

### 5.3 Evicting stale evidence from the 3D anchor reduces identity switches

Figure 2a shows the 3D anchor and the two correspondence terms it is scored with. Identity switches are the tracker’s main remaining association error, and stale evidence in the 3D anchor is one of their causes. When an identity’s 3D anchor retains detections without a limit, an identity that lost its animal much earlier continues to compete for assignments at full strength from the last position at which it was seen. If another animal then passes through that position while no recent appearance cue is available, the two identities swap. We therefore evicted retained detections older than *N* frames before matching (Section 4.14). Supplementary Figure 4f shows a sweep of *N* on all 74 SLAP-2M sessions, with every other setting at its default. At *N* = 20, within-view identity switches fell from 3,753 to 3,333, 11% fewer, and mean cross-view IDF1 over the 42 multi-animal sessions rose from 0.911 to 0.932. Twenty sessions improved and 6 worsened (two-sided Wilcoxon signed-rank test, *P* = 0.011).

Every eviction window gave fewer switches than no eviction, with 3,500 at *N* = 1, 3,406 at *N* = 10, 3,333 at *N* = 20 and 3,390 at *N* = 30 (7, 9, 11 and 10% fewer). Mean cross-view IDF1 did not follow the switches at the shortest window. It was 0.894 at *N* = 1, below the 0.911 with no eviction, and then 0.918, 0.932 and 0.929 at *N* = 10, 20 and 30. *N* = 20 beat *N* = 1 at the session level (*P* = 0.002), and the differences from *N* = 10 and 30 were small (*P* = 0.73 and 0.50). We chose *N* = 20 as the default because it gave the fewest switches and the highest mean cross-view IDF1 in the sweep. Eviction does not help on every dataset. On MouseDyad-10M, no eviction gave 102 switches against 130 at *N* = 20, and the session-level comparison of cross-view IDF1 favored no eviction (11 sessions better without it and 4 worse, *P* = 0.030).

### 5.4 Cross-view identity from 3D geometry alone in social-DANNCE rats

The social-DANNCE datasets record groups of rats with six calibrated cameras [13]. social-DANNCE keeps track of each rat by marking it with small patches of food coloring, and its center-of-mass network is trained on labels ordered by color. We asked whether the LUC3D tracker can recover identity from 3D geometry alone without using those markings (Supplementary Section 5.5).

Cross-view IDF1 was near perfect in most sessions (Supplementary Figure 4e). The median was 1.0 in all three datasets, and 35 of the 41 sessions scored exactly 1.0. Five of 6 TRIADS sessions, 5 of 6 BEDDING sessions and 26 of 29 SCN2A sessions had no identity switch. Each of the five sessions with a switch had one swap. In four of them the two rats came within 6 mm of each other at the swap, where their 3D tracks nearly coincide. The fifth happened during a body-overlap contact at 45 mm. One swap reverted after a single frame. The other four lasted to the end of the session. IDF1 assigns each tracker identity to one rat for the whole session. A lasting swap therefore splits the session in two, and the shorter part counts as wrong. Swaps close to either end of a session cost little (0.970 and 0.972, both within about the first 3% of the session). Swaps nearer the middle cost more (0.792 in TRIADS at frame 41,298 of 60,000, and 0.648 in SCN2A at frame 31,312 of 89,000). The 36 sessions without a swap contain 68,560 frames in which two rats are within 50 mm of each other, so close contact alone does not seem to cause swaps. Scoring against randomized ground truth gives an IDF1 of 0.500, which is chance. Splicing the ground truth mid-session gives exactly 0.5. Running the same detections through a deliberately wrong calibration drops the IDF1 to 0.519.

### 5.5 social-DANNCE benchmark

We used three dataset families from the social-DANNCE deposit [13]. TRIADS has 6 sessions of three male rats at 60,000 frames each. BEDDING has 6 sessions of two female rats on bedding at 90,000 frames each. SCN2A has 29 sessions of two rats at 89,000 frames each, from the first and third rounds of a round-robin pairing (14 and 15 sessions). The second round had no files in the deposit and was not used. Every session was recorded by six calibrated cameras at 50 frames per second, and the s-DANNCE 3D poses use a 23-node skeleton. The 2D detections were made by projecting each session’s 3D poses into the six views through that session’s calibration, which reproduces the deposit’s stored 2D labels to machine precision. The order of the animals was then shufÒed independently in every frame and every view, and the true identities were kept in a separate file used only for scoring. The tracker ran at the default settings of Section 4.8. Cross-view IDF1 was computed by matching tracker identities to animals once per session with a Hungarian assignment. A switch was counted when an animal’s majority identity across views changed between frames. The distance at a swap is the closest approach of the two swapped animals within 50 frames of it. The three controls re-scored a session against randomized ground truth, spliced the ground truth mid-session, and ran the detections through a rotated calibration. These detections are reprojections of the published 3D poses, so they contain no detector error. The benchmark therefore tests identity recovery from geometry, not robustness to a detector’s mistakes.

### 5.6 Calibration benchmark

The SLAP-2M rig (Section 4.3) has eight cameras at a resolution of 1280 × 1024, and four of its calibration sessions were used. An 18-camera dataset was recorded on a separate rig at 1680 × 1200 with one calibration session of 1,800 frames. The MouseDyad-10M rig has five cameras at 1280× 1024, and all five of its calibration sessions were used. In every session a single 8 × 11 ChArUco board of 24.0 mm squares with 18.75 mm markers from the 4 × 4 (1000) dictionary was moved by hand through the working volume. The SLAP-2M and 18-camera datasets were used to select default parameters for calibrat3. The MouseDyad-10M dataset was intentionally held out for testing.

Both calibration tools ran on OpenCV 4.13.0. calibrat3 vendors OpenCV.js 4.13.0 and cannot be run on another, and OpenCV 5 moved the ChArUco corner origin by half a pixel in both axes. A calibration scored across the two OpenCV versions pays an undue penalty. aniposelib 0.8.0 fits two intrinsic parameters per camera (focal length and *k*_1_) and pins the principal point at the image center. Its detector covers every frame in which a camera sees the board. That is a median of 2,606 frames per camera on the 2022-10-07 SLAP-2M session and 623 on the 18-camera session, where calibrat3’s sampled subset gives 882 and 378, so Anipose fits from more frames in both. calibrat3 ran in headless Chromium through its own interface at a sampling target of 800 frames, which an integer frame stride turns into 867 to 1,003 frames per session. It fits six intrinsic parameters per camera: *f_x_*, *f_y_*, *c_x_*, *c_y_*, *k*_1_ and *k*_2_. On the 18-camera rig it was given Camera 6 as the reference camera, the only one co-visible with both halves of that rig.

Every calibration was scored by aniposelib, which triangulates each 3D point from all cameras that detected it and reprojects it into each of them. Points that failed to triangulate under any one calibration were dropped from all of them, so every calibration is scored on one shared set of observations. To separate detection quality from solver quality, one run passed calibrat3’s detections to aniposelib’s solver. The calibrat3 solver was also re-run on identical detections with one setting changed at a time to isolate the effect of the intrinsic model. A synthetic rig with known cameras and an exactly planar board was used to measure absolute accuracy in focal length and scale. Anipose’s default gives it the larger frame budget, so aniposelib was also re-solved on only the frames calibrat3 used. The settings changed are given in Supplementary Section 5.8. Calibration time was measured with each tool run on its own on a 64-core machine. Anipose ran one detection process per camera.

### 5.7 Two-dimensional pose estimation models

The 2D detections in SLAP-2M and MouseDyad-10M came from top-down SLEAP models which were subsequently used for LUC3D’s performance evaluation [3]. Each model pair has a centroid network that finds every animal in the full frame and a centered-instance network that places the 15 keypoints in a crop around each centroid. Both networks used the U-Net backbone anchored on the trunk keypoint. The SLAP-2M models were trained in SLEAP 1.2.8 from random initialization on 9,068 labeled frames holding 18,752 instances. The MouseDyad-10M models were trained in SLEAP 1.3.3 on 39,644 labeled frames holding 76,279 instances. In both cases 5% of the labeled frames were held out at random for validation. The SLAP-2M predictions were made at a peak threshold of 0.4, and the MouseDyad-10M predictions at a peak threshold of 0.25. These predictions were used for the LUC3D cross-view tracking and triangulation comparisons (Section 4.7).

The SLAP-2M centered-instance model predicts keypoints with a median error of 3.0 pixels from user labels (90th percentile 12.1 pixels) and the MouseDyad-10M model with a median error of 1.6 pixels (90th percentile 4.8 pixels). In both datasets the shoulders were the hardest body keypoints and the nose and tail base the easiest. The lower accuracy of the SLAP-2M model is consistent with its smaller training set and the wider range of strains, coat colors, bedding and levels of enrichment (Section 4.3).

### 5.8 Calibration accuracy compared with Anipose

calibrat3 has the lower median reprojection error on all three rigs (Supplementary Figure 2a to f). On the held-out MouseDyad-10M rig, which had no part in choosing calibrat3’s defaults, the median is 0.07 pixels and Anipose 0.09. On the two rigs that were used to choose the defaults the margins are larger. On the SLAP-2M rig the calibrat3 median is 0.10 pixels and Anipose 0.32. On the 18-camera rig calibrat3 is 0.21 pixels and Anipose 1.28. calibrat3 is lower at both the median and the 95th percentile in all 31 cameras of the three rigs. The calibrat3 solver was tested on identical detections with one setting changed at a time to examine differences to the intrinsic model (Supplementary Figure 2g). Replacing calibrat3’s six free intrinsic parameters per camera with aniposelib’s focal length and *k*_1_ moves the median from 0.212 to 0.868 pixels. Removing outlier rejection moves it to 0.252. Freezing the intrinsics gives 1.235. Removing the rigid-board term has no measurable effect. Scoring on only the 900 held-out frames that calibrat3 did not use changed every configuration by less than 0.004 pixels. calibrat3 is faster than Anipose on the MouseDyad-10M rig and slower on the other two rigs (Supplementary Figure 2h).

Both calibrations are scored on one shared set of corners, but each was fitted from its own detector’s frames, and the two schedules differ. aniposelib also roots its extrinsic chain at its first camera, whereas calibrat3 was given Camera 6. Neither difference explains the gap between the two tools. Restricting aniposelib to the frames calibrat3 used lowered its median by about 0.1 pixels on the 18-camera dataset. Giving it the same reference camera had no effect. calibrat3’s median remained less than half of aniposelib’s in both cases.

Board-corner reprojection error is the quantity both solvers minimize. More free parameters can lower it by absorbing board non-planarity, and it is insensitive to a global scale error. On the synthetic rig, calibrat3 recovers the focal length to 0.05% and Anipose to 2.20% (Supplementary Figure 2i). The reconstructed rig scale differs from ground truth by 0.03% for calibrat3 and 2.98% for Anipose. With no board error to absorb, the additional parameters improve the recovered intrinsics.

These measurements come from three rigs in one laboratory using one board design. aniposelib’s solver is non-deterministic. On one MouseDyad-10M session it failed to converge on three of six draws from byte-identical detections. The three converged draws agree to within 0.002 pixels, and the figure reports one of them. calibrat3 is deterministic so does not require multiple runs on the same data.

### 5.9 Hardware-synchronized acquisition and live calibration with Panopticon

On a nine-camera rig recorded through Panopticon (Supplementary Figure 1a), an infrared LED driven from the trigger board’s stimulation output was registered by all nine cameras at every transition, 542 to 2,118 times above the per-pixel background, so a stimulation event falls on the same frame index in every view (Supplementary Figure 1b, c). Over a 19-minute recording the inter-frame interval had a median of 9.997 ms with a jitter of 2.3 *µ*s (Supplementary Figure 1d). The cameras’ timestamp clocks ran 211 to 223 parts per million below nominal and differed from one another by up to 12 parts per million, a drift of about 14 ms over the recording (Supplementary Figure 1e), so frames are aligned across cameras by trigger ordinal rather than by timestamp.

On a calibration recorded with the live coverage graph, Panopticon’s solver reached a median reprojection error of 1.51 pixels and calibrat3 0.15 pixels on the same 758,665 corner observations (Supplementary Figure 1f), and the focal lengths recovered by the two solvers agree to a median of 2.1% (Supplementary Figure 1g). The at-rig solve confirms the recording and the camera geometry before animals are placed in the arena, and the final calibration should be solved from the saved video by calibrat3 (Supplementary Figure 2) or Anipose.

